# *In vivo* characterization and application of a novel potassium channel-based optogenetic silencer in the healthy and epileptic mouse hippocampus

**DOI:** 10.1101/2021.09.06.459077

**Authors:** P Kleis, E Paschen, U Häussler, YA Bernal Sierra, CA Haas

## Abstract

The performance of available optogenetic inhibitors remains insufficient due to low light sensitivity, short-lasting photocurrents, and unintended changes in ion distributions. To overcome these limitations, a novel potassium channel-based optogenetic silencer was developed and successfully applied in various *in vitro* and acute *in vivo* settings (Bernal Sierra et al., 2018). This tool, a two-component construct called PACK, comprises a photoactivated adenylyl cyclase (bPAC) and a cAMP-dependent potassium channel (SthK). Here, we examined the long-term inhibitory action and side effects of the PACK construct in healthy and epileptic adult male mice. We targeted hippocampal CA1 pyramidal cells using a viral vector and enabled illumination of these neurons via an implanted optic fiber. Local field potential (LFP) recordings from the CA1 of freely moving mice revealed significantly reduced neuronal activity during 50-minute intermittent illumination, especially in the beta and gamma frequency ranges. Adversely, PACK expression in healthy mice induced chronic astrogliosis, dispersion of pyramidal cells, and generalized seizures. These side effects were independent of the light application and were also present in mice expressing bPAC without the potassium channel. Additionally, light-activation of bPAC alone increased neuronal activity, presumably via enhanced cAMP signaling. In chronically epileptic mice, the dark activity of bPAC/PACK in CA1 prevented the spread of spontaneous epileptiform activity from the seizure focus to the contralateral bPAC/PACK-expressing hippocampus. Taken together, the PACK tool is a potent optogenetic inhibitor but requires refinement of its light-sensitive domain to avoid unexpected physiological changes.

**Significance statement:** Optogenetics allows precise manipulation of neuronal activity via genetically encoded light-sensitive proteins. Unfortunately, available optogenetic inhibitors are not suitable for prolonged use. The newly developed two-component potassium channel-based optogenetic inhibitor, PACK, has been identified as a potent silencer of neurons in various acute experiments. Here, we characterized the PACK construct in freely behaving healthy and epileptic mice. We targeted the PACK silencer specifically to CA1 pyramidal neurons, where illumination with short light pulses at low frequencies reliably reduced neuronal activity. In chronically epileptic mice, PACK prevented the spread of epileptiform activity from the seizure focus to the contralateral PACK-expressing hippocampus. The major disadvantage of the PACK silencer is its light-sensitive domain, the bPAC adenylyl cyclase, which may induce side effects.

## Introduction

Cell type-specific inhibition techniques are required in neuroscience to investigate the contribution of neuronal populations in physiological and pathophysiological processes. Optogenetic silencing takes advantage of genetically encoded light-sensitive proteins, allowing to ‘switch off’ neurons of interest with high temporal and spatial precision. Currently available optogenetic inhibitors such as inward-directed chloride pumps (e.g. halorhodopsins) and outward-directed proton pumps (e.g. archaerhodopsins) have several limitations. Namely, they require continuous high-intensity illumination, which can have unexpected excitatory outcomes due to tissue heating, abnormal ion distributions, and strong rebound responses (Wiegert et al., 2017; Owen et al., 2019). Even brief activation of halorhodopsin can change the intracellular chloride concentration, cause a positive shift in GABA_A_ receptor reversal potential, and decrease the action potential threshold, leading to elevated network excitability (Raimondo et al., 2012; Alfonsa et al., 2015; Sørensen et al., 2017).

Using potassium (K^+^) current as the hyperpolarizing factor would be a better approach since the resting state of neurons is based on K^+^ conductance and is thus more physiological than pumping chloride or protons against their electrochemical gradients. Several synthetic light-activated K^+^ channels have been engineered which, however, pose shortcomings such as the requirement of a chemical cofactor (Banghart et al., 2004; Janovjak et al., 2010), utilizing UV light (Kang et al., 2013) or very low photocurrents in mammalian cells (Cosentino et al., 2015). A newly developed K^+^ channel-based optogenetic silencer could potentially overcome these limitations (Beck et al., 2018; Bernal Sierra et al., 2018). The two-component silencer, named PACK, comprises a soluble photoactivated adenylyl cyclase from the *Beggiatoa* bacterium (bPAC; Stierl et al., 2011) and a cyclic nucleotide-gated potassium channel from *Spirochaeta thermophila* (SthK; Brams et al., 2014). The blue light receptor in bPAC activates the cyclase domain thus increasing cytosolic cyclic adenosine monophosphate (cAMP), which subsequently opens the co-expressed SthK channels. Benefits of the PACK silencer include robust expression in mammalian cells, signal amplification, and large long-lasting K^+^ currents. Previously, PACK has been shown to reliably inhibit neuronal firing in cell cultures, acute slice preparations, and anesthetized mice (Bernal Sierra et al., 2018). However, a long-term application in awake mice has so far not been tested. A prolonged precise inhibition technique would be valuable for investigating the contribution of specific cell populations in pathologies such as epilepsy.

Mesial temporal lobe epilepsy (MTLE), the most common type of acquired focal epilepsy in adults, is characterized by spontaneous hippocampal seizures, which are often pharmacoresistant (Engel J., 2001; Kwan and Sander, 2004). MTLE is usually described as a unilateral disease since the seizures arise in one hemisphere, ipsilateral to the pathological abnormality. However, in some patients and MTLE mouse models, epileptic activity propagates to the contralateral hippocampus (Gloor et al., 1993; Mintzer et al., 2004; Meier et al., 2007; Popovic et al., 2012; Paschen et al., 2020). To scrutinize the performance of PACK and investigate the contribution of the contralateral hippocampus in MTLE, we applied PACK-mediated inhibition in the well-established intrahippocampal kainate (ihpKA) mouse model. This model recapitulates the major pathological features of human MTLE, such as unilateral hippocampal sclerosis with cell loss and gliosis accompanied by subclinical spontaneous seizures (Bouilleret et al., 1999; Riban et al., 2002; Janz et al., 2017). Contralateral CA1 cells in the ihpKA mouse model exhibit elevated activity-related cytoskeleton (Arc) gene expression during the chronic phase of epilepsy (Janz et al., 2018), suggesting that these cells are involved in the contralateral epileptiform activity.

Here, we aimed to validate the inhibitory action of the PACK silencer in principal neurons of hippocampal CA1 in freely moving mice. We investigated the long-term histological and electrophysiological effects of the PACK construct and its components (bPAC, AAV9.CaMKII.mCherry) *in vivo*. We present evidence that PACK activation persistently reduces neuronal activity during illumination with a frequency as low as 0.1 Hz. Furthermore, we applied the PACK silencer in chronically epileptic mice, where PACK-expression in CA1, contralateral to the seizure focus, prevented seizure spread between hemispheres.

## Materials and Methods

### Animals

Experiments were performed in adult 10 to 21 week-old transgenic male mice (C57BL/6-Tg(Thy1-eGFP)-M-Line) (Feng et al., 2000). For this study, 54 mice were used, each representing an individual experiment. Mice were kept in a 12 h light/dark cycle at room temperature (RT) with food and water *ad libitum*. All animal procedures were carried out following the guidelines of the European Community’s Council Directive of 22 September 2010 (2010/63/EU) and were approved by the regional council (Regierungspräsidium Freiburg).

### Stereotaxic intrahippocampal injections

Intrahippocampal saline/kainate and recombinant adeno-associated virus (AAV) injections were performed in one surgery in deeply anesthetized mice (ketamine hydrochloride 100 mg/kg, xylazine 5 mg/kg, atropine 0.1 mg/kg body weight, i.p.) as described previously (Heinrich et al., 2006; Häussler et al., 2012; Janz et al., 2017). Using a Nanoject III (Drummond Scientific Company, Broomall, Pennsylvania, USA), 50 nL of 0.9% sterile saline or 15 mM kainate (Tocris, Bristol, UK) was stereotaxically injected into the left dorsal dentate gyrus (DG) at coordinates relative to bregma: anterioposterior (AP) = −2.0 mm, mediolateral (ML) = −1.5 mm, and relative to the cortical surface: dorsoventral (DV) = −1.5 mm. Following kainate injection, the occurrence of behavioral *status epilepticus* was verified by observation of mild convulsion, chewing, immobility, or rotations, as described before (Riban et al., 2002; Tulke et al., 2019). Ten mice died as a consequence of *status epilepticus* and further two were excluded due extreme hippocampal atrophy (Table 1).

**Table 1.** Experimental groups and sample sizes in LFP analysis. From the initial number of mice that entered the experiment (total injected), some died owing to intrahippocampal kainate injections or implantations, and some were excluded due to electrode/optic fiber positions not in CA1 or due to hippocampal atrophy. The sample size indicates the number of mice included in LFP analysis.

| Group | Total injected | Died due to surgery | Excluded | LFP analysis |
| --- | --- | --- | --- | --- |
| saline + PACK | 12 | 2 | 4 | 6 |
| saline + bPAC | 7 | 1 | 0 | 6 |
| saline + mCherry | 4 | 1 | 1 | 3 |
| saline | 7 | 0 | 0 | 7 |
| kainate + PACK | 11 | 4 | 2 | 5 |
| kainate + bPAC | 9 | 6 | 0 | 3 |
| kainate | 4 | 0 | 0 | 4 |

For optogenetic inhibition of CA1 principal cells, an AAV carrying the PACK silencer with a red fluorescent marker mCherry under the control of the Ca^2+^/calmodulin-dependent kinase II alpha (CaMKIIα) promoter (AAV9.CamKIIα:SthK-P2A-bPAC-mCherry referred to as AAV9.CaMKIIα:PACK-mCherry) was injected into the right hippocampus. In order to attribute the effects of AAV9.CaMKIIα:PACK-mCherry to its components, we included three control groups, in which mice were injected with (1) a construct carrying the adenylyl cyclase bPAC but lacking the SthK channel (AAV9.CamKIIα:bPAC-mCherry), (2) a construct carrying only mCherry (AAV9.CamKIIα:mCherry), or (3) no virus (for sample sizes, see Table 1). The viral constructs were injected into the right dorsal CA1 area (AP = -2.0 mm, ML = -1.3 mm, DV = -1.15 mm). The volume of the injected virus was adjusted to achieve an optimal and comparable expression pattern in CA1: (1) PACK – 300 nL, (2) bPAC – 250-300 nL, (3) mCherry – 300 nL with 1:5 dilution. All viral constructs were obtained from the Viral Core Facility, Charité – Universitätsmedizin Berlin, Germany.

### Electrode and optic fiber implantations

Implantations were performed 14-19 days after intrahippocampal injections as described previously by Janz et al. (2018). For local field potential (LFP) analysis, we implanted a Teflon-coated platinum-iridium wire electrode (125 µm diameter; World Precision Instruments, Sarasota, Florida, USA) into each hippocampus: AAV-injected right CA1 (AP = −2.0 mm, ML = -1.4 mm, DV = −1.1 mm) and saline/kainate-injected left DG (AP = −2.0 mm, ML = +1.4 mm, DV = −1.6 mm). An optic fiber (ferrule 1.25 mm, cannula 200 µm diameters; Thorlabs Inc., Newton, New Jersey, USA) was implanted into the AAV-injected CA1, adjacent to the electrode at a 30° angle (AP = −2.0 mm, ML = -2.4 mm, DV = −1.0 mm). Two stainless steel screws (DIN 84; Schrauben-Jäger, Landsberg, Germany) were implanted above the frontal cortex to provide a reference and ground. Electrodes and screws were soldered to a micro-connector (BLR1-type) and fixed with dental cement (Paladur, Kulzer GmbH, Hanau, Germany). The electrode and optic fiber positions were confirmed by post hoc histology (Fig.1-1). Five mice were excluded from LFP analysis due to electrode/optic fiber locations in the cortex above CA1. Three mice died following implantation procedures (Table 1).

**Figure 1.**
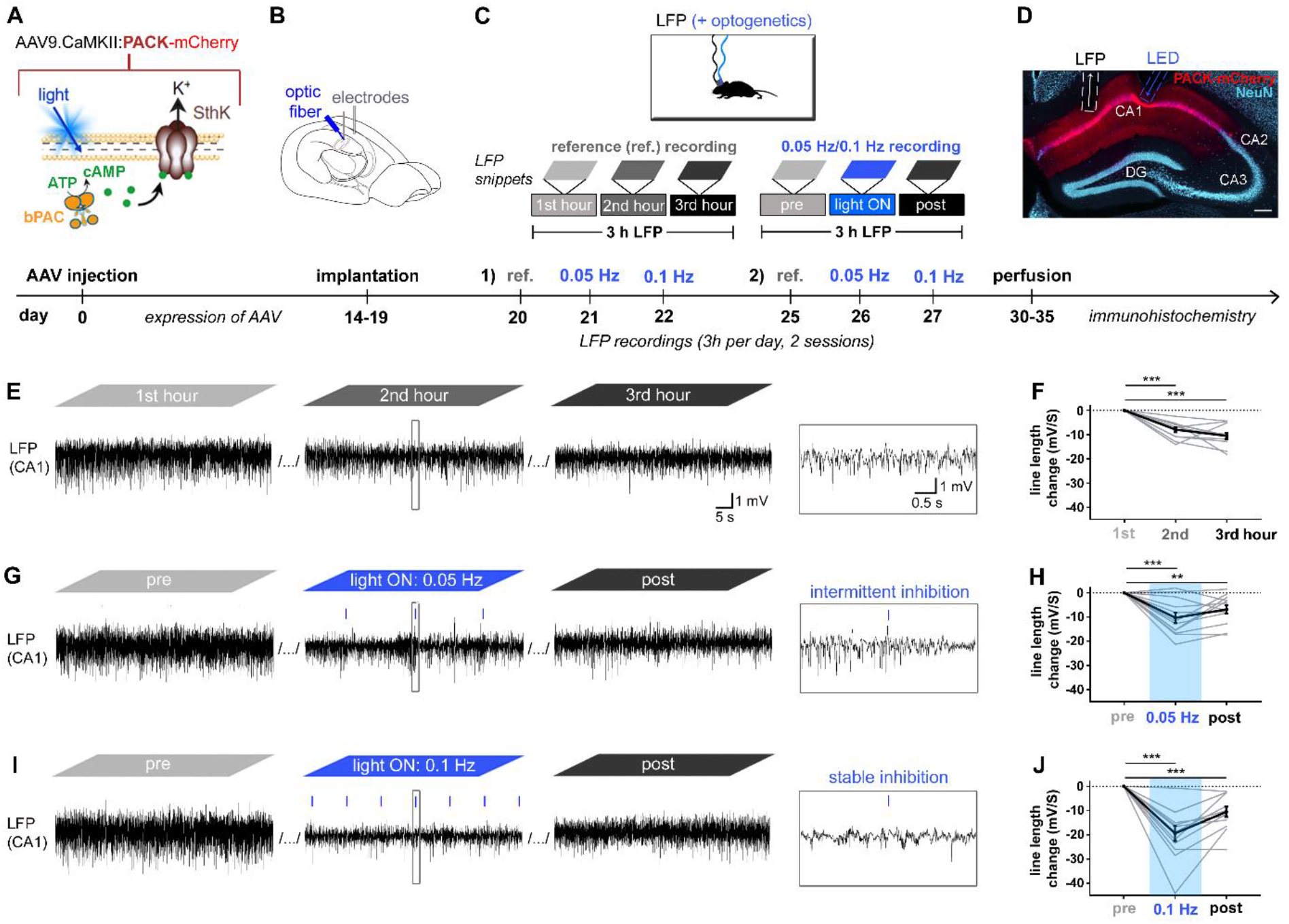
PACK-mediated optogenetic inhibition of CA1 principal cells *in vivo*. (A-D) Experimental design with a timeline. (A) We targeted PACK (bPAC+Sthk) to excitatory neurons by injecting the AAV9.CaMKII vector into the CA1 region of the hippocampus. (B) After implantations of wire electrodes and optic fibers into CA1, (C) we recorded LFPs in freely moving mice for three hours a day. Reference (ref.) recordings were without illumination, whereas the ‘0.05 Hz/0.1 Hz recordings’ included one-hour pre-recording, light ON phase with 0.05 Hz or 0.1 Hz illumination, and a post-recording. Each recording type (ref., 0.05 Hz, 0.1 Hz) was performed twice in each mouse (for details see Materials and Methods). (D) PACK-mCherry expression and electrode/optic fiber positions in CA1 were confirmed by histology at the end of the experiment. (E, G, I) Representative LFP snippets from the first, second, and third hour of each recording type. (E) In ref. recordings, LFP magnitude decreased over time, (F) confirmed by the significant drop of mean line length (in black) in the second and third recording hour. (G) Applying 10 ms blue light pulses (∼80 mW/mm^2^, 460 nm) at 0.05 Hz resulted in an intermittent reduction of LFP magnitude. (H) Mean line length was significantly reduced during 0.05 Hz light application and in the post-recording. (I) Applying light pulses at 0.1 Hz resulted in a stable reduction of LFP magnitude, (J) which is reflected in a robust reduction of line length during 0.1 Hz illumination. One-sample t-test (n=12 trials from 6 mice, grey lines), ** p<0.01, ***p<0.001. Mean presented in black with SEM as error bars.

### Electrophysiological recordings and optogenetic manipulations

Three-hour-long LFPs were acquired from freely moving mice in the period of 19-40 days after intrahippocampal injections. For LFP recordings, mice were connected to a miniature preamplifier (MPA8i, Smart Ephys/ Multi Channel Systems, Reutlingen, Germany). Signals were amplified 1000-fold, bandpass-filtered from 1 Hz to 5 kHz and digitized with a sampling rate of 10 kHz (Power1401 analog-to-digital converter, Spike2 software, Cambridge Electronic Design, Cambridge, UK).

For each AAV9-injected mouse (with PACK, bPAC or mCherry), we acquired reference LFPs before, in between, and after illumination experiments. LFPs with one-hour illumination at 0.05 Hz or 0.1 Hz were recorded twice per frequency on separate days. Each mouse represents a biological replicate (n=3-7 per group, Table 1) and the number of recordings per mouse a technical replicate (n=2 per recording type). The sessions with illumination comprised one-hour pre-recording, ‘light ON’ phase during the second hour and post-recording during the third hour. During the ‘light ON’ phase, blue light pulses (460 nm, ∼80 mW/mm^2^, 10 ms pulse duration, blue LED from Prizmatix Ltd. Givat-Shmuel, Israel) were applied every 10 or 20 seconds. To hinder rebound excitation resulting from the illumination off-set (Mahn et al., 2016; Bernal Sierra et al., 2018), each light pulse had a 5 ms ramp-like termination (within pulse fade-off). Furthermore, during the last 10 minutes of the ‘light ON’ phase, 5 ms light pulses were applied with gradually reducing intensity (within recording fade-off). The light pulse duration and frequencies were selected based on previous work by Bernal Sierra et al. (2018), who demonstrated that shining 5 ms light pulses with 0.05 Hz in hippocampal slices provided long-lasting stable inhibition of current-elicited spiking in PACK-expressing CA1 pyramidal cells.

Kainate-injected mice, which were implanted with intrahippocampal electrodes but did not receive a viral vector or an optic fiber, served as epileptic ‘no virus’ controls. Three-hour recordings from these mice were performed on day 35 or 36 after kainate injection.

### Analysis of local field potentials

LFP data were visually inspected with Spike2 software and analyzed in detail using Python 2.7. The line length, a sum of distances between successive data points, was selected as a measure of the LFP waveform dimensionality since it is sensitive to variations in both amplitude and frequency (Esteller et al., 2004). We calculated the line length of downsampled data (500 Hz) by using the following equation, where *L* is the line length (mV/s), *x* is the data-trace, *k* is one data point, and *t* is the recording duration in seconds:

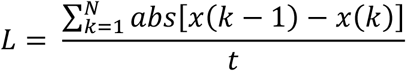

The line lengths were calculated for the first, second, and third hour of each recording. In recordings with illumination, the line length calculations and spectral analysis were done on LFP data recorded during the first 50 minutes; the last 10 minutes were omitted due to light intensity fade-off. Furthermore, in recordings with illumination, the line lengths shortly after the light pulses were compared to the line lengths directly before the light pulses. For this, two-second snippets were extracted before and after each light pulse during the first 50 minutes of the ‘light ON’ phase. In addition, two-second snippets at corresponding time points were extracted in pre-recordings. Subsequently, the mean “pre”, “before pulse”, and “after pulse” line lengths were calculated for each recording session (2 per animal). Periods with electrographic generalized seizures (GS), where hypersynchronous neuronal activity was propagating across hemispheres followed by postictal depression, were removed from analysis due to strongly altered LFP characteristics.

The frequency compositions of the LFPs are presented with power spectral densities (PSD) and spectrograms generated by fast Fourier transform (FFT) of LFP raw data (sampling rate 10 kHz). The PSDs were estimated by applying Welch’s method in Python 2.7 (scipy.signal.welch function), based on time averaging over short periodograms (periodogram length = 10x sampling rate). The oscillatory power was calculated as the area under the PSD plot for the respective frequency ranges: delta (1-4 Hz), theta (4-12 Hz), beta (12-30 Hz), gamma (30-120 Hz). For time-frequency representation of the LFP power in a spectrogram, the Hanning window was applied and data in the time-domain (length of FFT= 10x sampling rate) was broken up into overlapping (overlap = 0.25x sampling rate) segments (segment length = 1x sampling rate).

### Analysis of epileptiform activity

Downsampled hippocampal LFPs recorded from epileptic animals were analyzed in detail using a custom-made semi-automated algorithm that detects and classifies epileptiform activity (Heining et al., 2019). In the ihpKA mouse model, epileptiform activity occurs as single sharp wave epileptiform spikes and as bursts, which are clusters of many spikes (Riban et al., 2002). The algorithm classifies the bursts according to their spike-load into low-load, medium-load, and high-load bursts as described by Heining et al. (2019). To assess the effect of PACK-mediated inhibition on seizure activity, we calculated the ‘burst ratio’, which is the duration of high-load bursts per total recoding time. The automatic detection of high-load bursts was verified by visual inspection of the LFP recordings. Sessions with GS were removed from the analysis due to long-lasting suppression of neuronal activity after such a seizure.

### Perfusion and tissue preparation

After the last LFP recording (at day 30-40), the mice were deeply anesthetized and transcardially perfused with 0.9% saline followed by 4% paraformaldehyde in 0.1 M phosphate buffer (PB, pH 7.4). The brains were dissected and post-fixated overnight in 4% paraformaldehyde, followed by sectioning (coronal plane, 50 μm) with a vibratome (VT100S, Leica Biosystems, Wetzlar, Germany). The slices were collected and stored in PB for immunohistochemistry.

### Immunohistochemistry

To determine optic fiber and electrode positions, to visualize the hippocampal anatomy after AAV9-injections, and to validate hippocampal sclerosis in ihpKA mice, we performed immunohistochemistry with markers for neurons (NeuN) and astrocytes (glial fibrillary acidic protein – GFAP). For the immunofluorescence staining, free-floating sections were pre-treated with 0.25% TritonX-100 and 10% normal horse or goat serum (Vectorlabs, Burlingame, California, USA) diluted in PB for 1 h. Subsequently, slices were incubated either with guinea-pig anti-NeuN (1:500; Synaptic Systems, Göttingen, Germany) or rabbit anti-GFAP (1:500, Dako, Glostrup, Denmark) overnight at 4°C. Sections were rinsed and then incubated for 2.5 h in donkey anti-guinea-pig or donkey anti-rabbit Cy5-conjugated secondary antibody (1:200, Jackson ImmunoResearch Laboratories Inc., West Grove, Pennsylvania, USA), followed by extensive rinsing in PB. The sections were mounted on glass slides and coverslipped with Immu-Mount^™^ mounting medium (Thermo Shandon Ltd, Runcorn, UK).

### Image acquisition and histological analysis

Tiled fluorescent images of the brain sections were taken with an *AxioImager 2* microscope (Carl Zeiss Microscopy GmbH, Jena, Germany) using a Plan-Apochromat 10x objective with numerical aperture 0.45 (Zeiss, Göttingen, Germany). The exposure times (Cy5-labeled NeuN: 5 s, Cy5-labeled GFAP: 3 s, mCherry: 300 ms) were kept constant for each staining to allow for comparisons across animals.

To assess the effect of AAV9-mediated delivery of PACK, bPAC and/or mCherry and long-term expression of these proteins on hippocampal histology, we measured the relative expression intensities of GFAP labeling in the mCherry-expressing dorsal CA1 at three positions along the anteroposterior axis (−1.70 mm, -1.94 mm, and -2.18 mm from bregma). The quantification was performed in Fiji ImageJ by drawing a polygon-shaped area of interest (ROI) around mCherry expression in CA1 (*str. oriens* to *str. radiatum*) and taking the mean grey area of both mCherry and the GFAP labeling within this ROI. Areas with glial scarring around the implantations were excluded by adjusting the ROI. The same measurement was done in the contralateral CA1 by drawing a similar ROI, which avoided implant-related scars. For normalization, in each slice the local background was measured in a small square (41457 µm^2^) in the cortex and subtracted from the mean grey areas of each ROI in CA1. Finally, mean expression intensities of mCherry and GFAP in left (saline-injected) and right (virus-injected) CA1 were presented for each animal. Furthermore, we quantified the width of the pyramidal cell layer in dorsal medial CA1 by measuring three perpendicular lines in the left and in the right CA1 of each NeuN-labeled section (3 sections per mouse, AP: -1.70 mm, - 1.94 mm, -2.18 mm from bregma) and compared the mean width of the left and right CA1 pyramidal cell layer.

In ihpKA mice, the presence of hippocampal sclerosis in the kainate-injected hippocampus was confirmed in NeuN-labeled sections showing granule cell dispersion and cell loss in CA1 and CA3 and in GFAP-labeled sections demonstrating astrogliosis.

### Statistical analysis

Data were tested for statistical significance with GraphPad Prism 8 software (GraphPad Software Inc.). To determine how the line length and spectral power changed during the second and third hour of LFP recordings, the baseline value (in the first hour) was subtracted from the original values and tested for significance using a one-sample t-test with a Bonferroni correction of the significance level (α=0.025 for two comparisons). A paired t-test was used for comparing two matched groups of parametric data. Comparisons of more than two parametric data sets were performed either with a one-way ANOVA or with repeated-measures (RM) ANOVA, in case of matched groups. If an ANOVA indicated that not all group means were equal, Tukey’s multiple comparisons test was performed additionally. Friedman’s test (matched) or Kruskal-Wallis test (non-matched) with Dunn’s post hoc were applied for comparing three groups of non-parametric data. Pearson’s correlation coefficient was used to measure the strength of association between two variables. Significance thresholds were set to: *p<0.05, **p<0.01 and ***p<0.001. For all parametric data, mean and SEM are given; for non-parametric data, median with interquartile range (IQR) are reported.

## Results

### Light-activated PACK reduces the activity of pyramidal cells *in vivo*

To verify the inhibitory action of the PACK silencer in awake mice, we targeted hippocampal principal cells by locally injecting AAV9.CaMKIIα.PACK-mCherry into the CA1 area of the dorsal hippocampus (Fig. 1A) and enabled illumination onto these neurons via an implanted optic fiber (Fig. 1B). To test whether applying short light pulses (10 ms) at low frequencies *in vivo* results in sustained inhibition of PACK-expressing CA1 neurons as previously demonstrated *in vitro* (Bernal Sierra et al., 2018), we shined blue light at 0.05 Hz and 0.1 Hz for one hour. The light ON phase was enclosed by a pre- and post-recording, one hour each (Fig. 1C). Following the recording phase, histological analysis revealed that expression of PACK-mCherry was restricted to pyramidal neurons in CA1 with labeling in cell bodies and dendrites (Fig.1D). For LFP analysis, we only included mice, which had the optic fiber and recording electrode positioned in CA1 (Supplementary Fig. 1, n=6).

First, we recorded three-hour reference LFPs in each mouse (Fig. 1E) to control for the change in LFP characteristics occurring without any manipulation due to habituation - reduction in arousal and exploratory behavior (Cabib et al., 1990; Hinman et al., 2011). Quantification of the LFP signal by determination of the line length, a measure for LFP magnitude (see Materials and Methods), revealed an innate decrease during the reference recordings (first hour: set to zero; second hour: -7.98 ± 1.01 mV/s, one-sample t-test: t=7.93, n=12 trials, p<0.0001, α=0.025; third hour: -10.54 ± 4.35 mV/s, one-sample t-test: t=8.38, n=12 trials, p<0.0001, α=0.025; Fig. 1F). Next, we activated PACK with intermittent light application. During 0.05 Hz illumination, LFP magnitude was reduced directly after light pulses, followed by periods of recovery (Fig. 1G). The mean line length significantly decreased during 0.05 Hz illumination (50 minutes) compared to the pre-recording (−10.33 ± 2.18 mV/s, one-sample t-test, t=4.73, n=12 trials, p=0.0006, α=0.025; Fig. 1H). Illumination with 0.1 Hz provided a stable reduction of the LFP magnitude (Fig. 1G) with a strong decrease in the line length during the light ON phase (−19.42 ± 3.14 mV/s, one-sample t-test: t=6.18, n=12 trials, p<0.0001, α=0.025; Fig. 1J). The change of line length from the first to the second hour was significantly different in the three recording types (reference: -7.98 ± 1.01 mV/s, 0.05 Hz: -10.33 ± 2.18 mV/s, 0.1 Hz: -19.42 ± 3.14 mV/s, RM ANOVA: F=14.40, n=12, p= 0.012). The drop of line length was notably higher in the 0.1 Hz recordings than in the respective reference recordings (Tukey’s multiple comparison test: p=0.003), whereas in the sessions with 0.05 Hz it was similar to the reference recordings (Tukey’s multiple comparison test: p=0.33).

To test the reliability of the PACK-mediated inhibition, we further analyzed the responses to light pulses applied at 0.05 Hz and 0.1 Hz. We extracted two-second LFP snippets before and after each light pulse (Fig. 2A), plotted their overlay (Fig. 2B, D), and calculated the mean of the “before pulse” and “after pulse” line lengths for each recording session (Fig. 2C, E). We also extracted two-second LFP snippets at corresponding time points from respective pre-recordings (“pre”) to serve as a baseline. In the 0.05 Hz session, reduction of LFP magnitude was reliable and reversible since there was a reduction after each light pulse, followed by a complete recovery during the 20-second interval between subsequent pulses (Fig. 2B). The mean “after pulse” line length was significantly smaller than “pre” and “before pulse” line length in 0.05 Hz sessions (“pre”: 48.67 ± 3.83 mV/s, “before pulse”: 48.38 ± 4.02 mV/s, “after pulse”: 32.39 ± 2.31 mV/s, RM ANOVA: F=17.83, n=12, p=0.0001, Tukey’s multiple comparisons test: p<0.001; Fig. 2C). Shining 10 ms light pulses every 10 seconds provided a persistent reduction in LFP amplitude (Fig. 2D). In the 0.1 Hz sessions, the line length was lower during the whole light ON period, including “after pulse” and “before pulse” snippets, suggesting a constant inhibitory effect (“pre”: 49.99 ± 4.80 mV/s, “before pulse”: 35.48 ± 3.13 mV/s, “after pulse”: 28.69 ± 2.23 mV/s, RM ANOVA: F= 18.02, n=12, p=0.0019, Tukey’s multiple comparisons test: “pre” vs. “before pulse” p= 0.0259, “pre” vs. “after pulse” p=0.003; Fig. 2E). In summary, shining blue light at 0.1 Hz onto PACK-expressing CA1 neurons *in vivo* resulted in a sustained reduction of the net neuronal activity throughout the light ON period.

**Figure 2.**
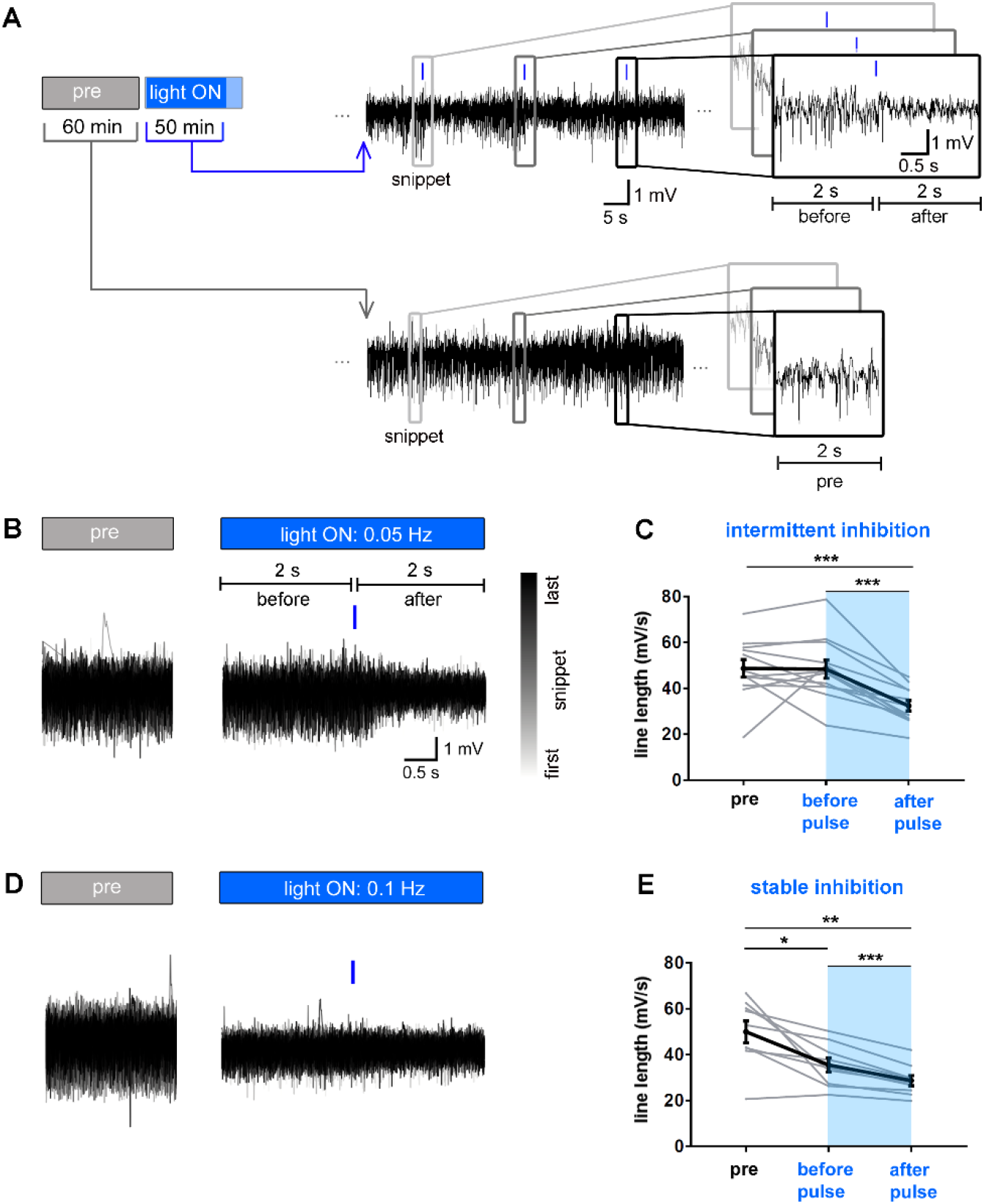
PACK-mediated inhibition *in vivo* is reliable and reversible. (A) Analysis of responses to 10 ms blue light pulses during the first 50 minutes of the light ON phase. Two seconds long LFP snippets before and after each light pulse were extracted and overlaid with a color-coding from grey to black (first to last LFP snippet). To enable comparison to baseline two-second snippets at corresponding time points were extracted from pre-recordings. (B, D) Representative plots of overlaid LFP snippets from a pre-recording and light ON recording with (B) 0.05 Hz and (D) 0.1 Hz illumination. (C, E) Line lengths (mV/s) were calculated for each two-second LFP snippet and the mean “pre”, “before pulse”, and “after pulse” line length for each recording (n=12) is presented (in grey). The mean (in black) “after pulse” line length was significantly smaller than “pre” and “before pulse” line length in both illumination modes, indicating a reliable reduction of neuronal activity. (C) Baseline (“pre”) and “before pulse” line lengths were similar in 0.05 Hz sessions, demonstrating recovery from inhibition before the next pulse was applied. (E) LFP line length was reduced throughout the 0.1 Hz light ON period (in blue), including before and after pulse phases, suggesting the inhibition was stable. RM ANOVA and Tukey’s multiple comparisons test (n=12 trials from 6 mice, grey lines), *p<0.05, ** p<0.01, ***p<0.001. Mean presented in black with SEM as error bars.

Next, we examined how light-induced PACK-activation in principal cells alters oscillatory activity in CA1. The most dominant oscillations measured in the hippocampus of freely behaving rodents are theta (4-10 Hz) and gamma (30-120 Hz) waves (Bragin et al., 1995; Buzsáki et al., 2003; Colgin, 2016). The theta and gamma peaks are prominent in spectrogram snippets (Fig. 3A) and in the mean PSD plot (PSD averaged across recordings; Fig. 3B) of the reference recordings acquired from PACK mice. We quantified the oscillatory power by taking the area under the curve of the PSD plot in the respective frequency range. The power of theta, beta, and gamma oscillations decreased significantly from the first to second hour presumably due to habituation (theta: -0.12 ± 0.02, beta: -0.13 ± 0.02, gamma: -0.16 ± 0.03, one-sample t-test, n=12, p: <0.0001-0.003, α=0.006; Fig. 3E)

**Figure 3.**
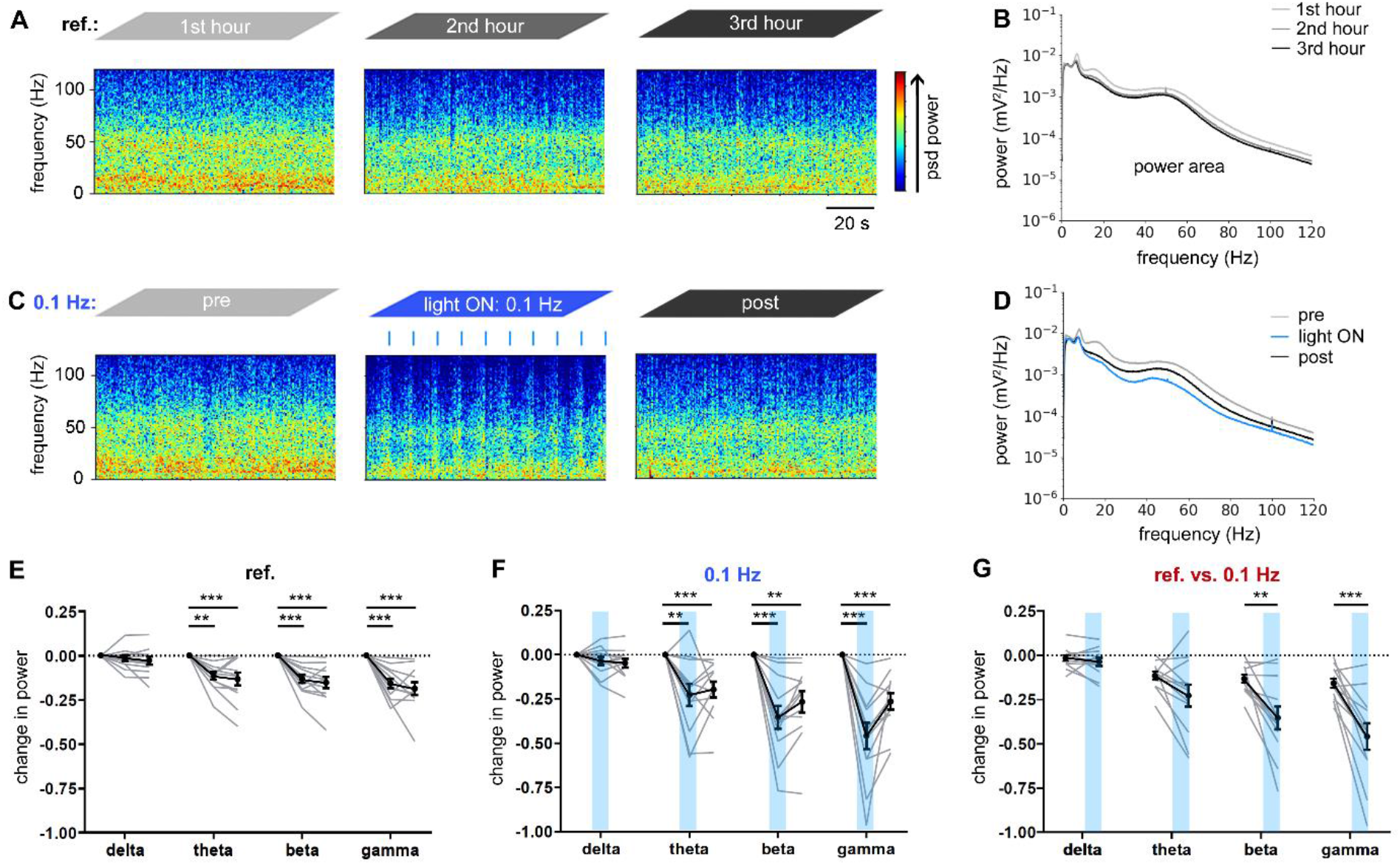
Spectral analysis reveals reduced power of beta and gamma oscillations during light-activation of PACK. (A) Representative spectrogram snippets from a ref. recording show baseline activity; blue colors reflecting low power and red colors reflecting high power of the corresponding frequencies. (B) Mean power spectral density (PSD) during the first (light grey), second (dark grey), and third (black) recording hour. (C) Light pulses applied at 0.1 Hz transiently decreased the spectral power. (D) The mean PSDs from pre-(light grey), light ON (blue), and post-recordings (black). (E-G) The oscillatory power was calculated by taking the AUC in the PSD plot for the particular frequency range. The power change is presented as baseline corrected values for the second and third hour. During the(E) ref. and (F) illumination recordings with 0.1 Hz, the power of theta, beta, and gamma declined significantly. (G) Change in power in the second hour of ref. versus 0.1 Hz recordings revealed that only beta and gamma power were further reduced by the light-induced PACK activation.

Applying light pulses at 0.1 Hz, transiently reduced the spectral power of frequencies above ∼10 Hz (Fig. 3C). The mean PSD during the 50-minute 0.1 Hz light ON phase was visibly reduced compared to pre- and post-recordings, especially in the beta and gamma ranges (Fig. 3D). Quantification of the power change revealed a significant drop of theta, beta, and gamma power during 0.1 Hz illumination (theta: -0.23± 0.06, beta: -0.35 ± 0.07, gamma: -0.46 ± 0.07, one-sample t-test, p<0.0001-0.0038, α=0.006, n=12 trials; Fig. 3F). The reduction of beta and gamma power during 0.1 Hz illumination was significantly stronger than the inherent decline in respective reference recordings (beta: paired t-test, t=3.63, n=12 trials, p=0.004, gamma: paired t-test, t=4.47, p=0.001, α=0.0125; Fig. 3G). These data indicate that PACK-mediated inhibition of CA1 neurons *in vivo* alters the power of beta and gamma oscillations.

### Light-dependent hyperactivity in bPAC-expressing mice

Activation of the PACK silencer includes cAMP production by soluble bPAC, which then opens the co-expressed SthK potassium channels in the cell membrane. The second messenger molecule, cAMP, is an important component of intracellular signaling, regulating the plasticity and excitability of neurons (Huang et al., 1994; Weisskopf et al., 1994; Gruart et al., 2012; Fukaya et al., 2021). Therefore, it is crucial to investigate whether activation of bPAC alone affects network excitability.

To this end, we targeted bPAC with the AAV9.CaMKII viral vector to CA1 neurons and repeated the experiments like with PACK mice (Fig. 4A-D). Surprisingly, light-activation of bPAC at 0.1 Hz led to sustained neuronal hyperactivity that is clearly visible in LFP snippets (Fig. 4E) as well as in corresponding spectrograms (Fig. 4G). The line length was significantly increased during 0.1 Hz illumination (12.10 ± 3.55, one-sample t-test: t=3.41, n=12, p=0.0058, α=0.025; Fig. 4F). The oscillations, which were mainly altered by bPAC activation were beta and gamma with significantly elevated power during illumination (beta: 0.50 ± 0.17, one-sample t-test: t=3.03, n=12, p=0.011, gamma: 0.34 ± 0.096, one-sample t-test: t=3.52, n=12, p=0.0048, α=0.0125; Fig. 4H-I). The theta oscillatory power increased slightly but not significantly during light-induced bPAC activation (0.25 ± 0.13, one-sample t-test: t=1.91, n=12, p= 0.08, α=0.0125; Fig. 4I). In the post-recording, LFP magnitude and gamma power dropped again, suggesting that the neuronal hyperactivity was reversible and related to light-induced elevation of cAMP levels (line length: -8.86 ± 2.80 mV/s, one-way t-test, t=3.16, n=12, p=0.0091, α=0.025; Fig. 4F; gamma power: -0.11 ± 0.04 mV/s, one-way t-test, t=2.68, n=12, p= 0.02; Fig. 4I). The elevation in neuronal activity was not induced by blue light *per se* since mice injected with AAV9.CaMKII.mCherry did not show any changes in the LFP signal during illumination (Supplementary Fig. 2). In summary, light-induced activation of bPAC without the SthK channel co-expression transiently elevates neuronal activity in CA1.

**Figure 4.**
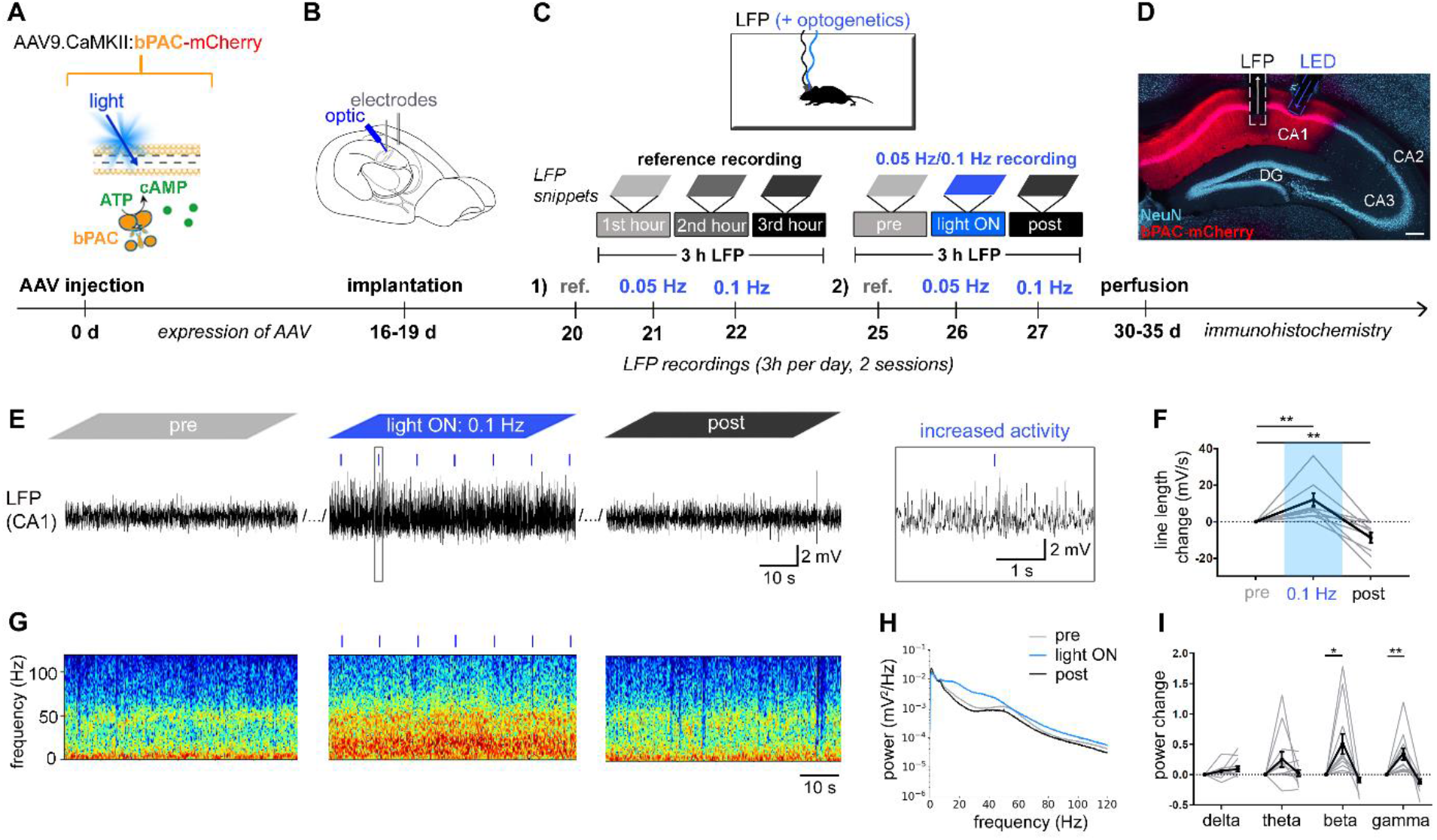
Light-activation of bPAC reversibly increases neuronal activity in CA1. (A) Experimental design. We targeted bPAC-mCherry to excitatory neurons in the CA1 area using the AAV9 vector. (B) Implantation, (C) recordings and (D) histological analysis were performed like in PACK mice. Scale bar 200 µm. (E) Representative snippets of a 0.1 Hz recording show persistently increased LFP magnitude during the light ON phase in bPAC-expressing CA1. (F) In the light ON phase, the mean line length was significantly increased compared to the baseline (pre) level. In the post-recordings, the line length was reduced compared to the baseline. (G) The spectrograms, taken from the same time window as LFP snippets, demonstrate elevated spectral power. (H) The mean PSD during 0.1 Hz illumination was increased at frequencies above ∼10 Hz. (I) The beta and gamma powers were significantly higher during the light ON phase compared to the pre-level. One-sample t-test (n=12 trials from 6 mice, grey lines), *p<0.05, ** p<0.01. Mean presented in black with SEM as error bars.

### Spontaneous generalized seizures arising in PACK- and bPAC-expressing mice

In healthy control mice, which received intrahippocampal saline and recording electrodes, epileptiform activity is normally absent in the LFP (Twele et al., 2017). Unexpectedly, in the majority of the PACK (5 out of 6) and all of the bPAC mice (n=6), hypersynchronous activity, spreading across both hemispheres, arose at least once during LFP recordings (Fig. 5A, B, E). Most of these electrographic generalized seizures were accompanied by behavioral correlates such as freezing, nodding, forelimb clonus, or rearing according to the Racine scale (Racine, 1972).

**Figure 5.**
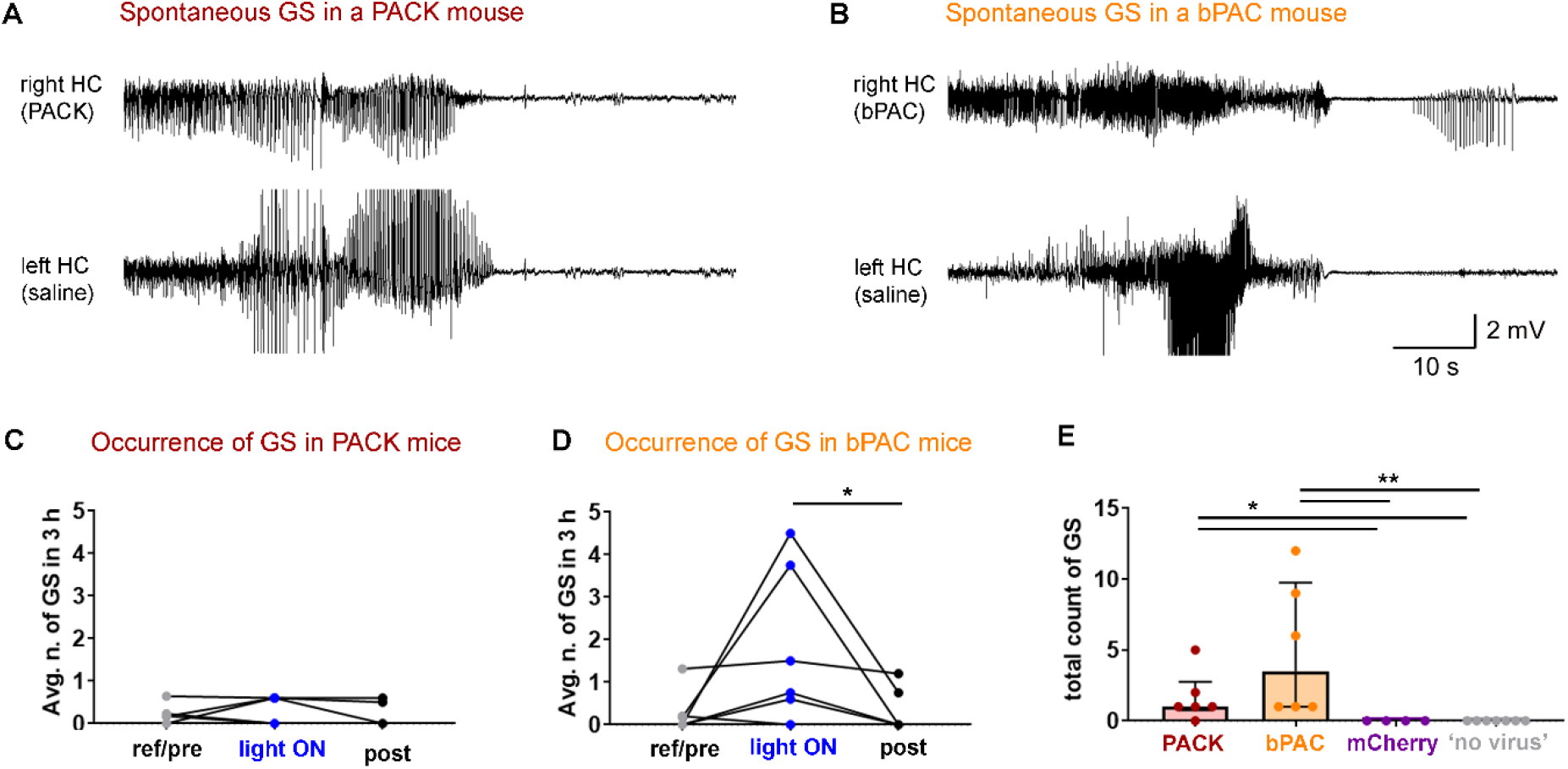
Spontaneous generalized seizures (GS) in PACK and bPAC mice during LFP recordings. (A) Representative spontaneous electrographic GS in a PACK mouse. The hypersynchronous activity spread across both hemispheres and was followed by postictal depression. (B) Similar GS were recorded in bPAC mice. (C,D) The number of GS was counted in all LFP recording types (baseline: “ref/pre”, “light ON”, “post”) of PACK (n=6) and bPAC (n=6) mice, and the average number of GS in three hours is presented and compared with Friedman’s test followed by Dunn’s multiple comparisons. The occurrence of GS in baseline recordings indicates seizure generalization independent from light-induced PACK/bPAC activation. (E) The total count of GS in LFP recordings (9-28 h) was the highest in bPAC mice. Control mice that expressed either mCherry in CA1 or no virus, experienced no seizure-like activity during recordings. Kruskal-Wallis test with Dunn’s multiple comparisons, * p<0.05, ** p<0.05. Median presented with IQR as error bars.

The occurrence of generalized seizures in baseline recordings (“ref” and “pre”) indicates seizure initiation independent from light-induced PACK (Fig. 5C) and bPAC (Fig. 5D) activation. Illumination of bPAC-expressing neurons increased the median of average seizure count in three hours compared to post-recordings (“pre”: 0.09 [0, 0.48], “light ON”: 1.13 [0.45, 3.94], “post”: 0 [0, 0.86], Friedman test: Fr= 6.38, n=6, p=0.041, Dunn’s post hoc: p=0.042; Fig. 5D). The median number of generalized seizures in all recordings was the highest in bPAC mice (3.50 [1.00, 9.75]; Fig. 5E). Most PACK mice experienced one generalized seizure throughout all the recordings (in total 9-28h), (1.00 [0.75, 2.75], n=6), whereas mCherry (n=4) and ‘no virus’ (n=7) control mice had no generalized seizures (Fig. 5E). These results suggest that the dark activity of bPAC is responsible for spontaneous generalized seizures arising in mice that express PACK or bPAC in CA1 pyramidal cells.

### Histological abnormalities in PACK- and bPAC-expressing CA1

An optogenetic tool suitable for long-term *in vivo* experiments should preserve the normal physiology and histology in the target area. For histological analysis, PACK, bPAC, and mCherry mice were perfused after the last LFP recording, 30-35 days after the intrahippocampal virus and saline injections. Coronal sections were immunolabeled with anti-NeuN and anti-GFAP to investigate the histology of neurons and astrocytes respectively.

To our surprise, we found notable widening of the pyramidal cell layer in PACK-expressing CA1 (Fig. 6A). The mean width of the pyramidal cell layer in PACK-expressing right CA1 was significantly higher than in the saline-injected left side (right CA1: 71.66 ± 2.01 µm; left CA1: 60.68 ± 0.54 µm, paired t-test: t=5.25, n=8, p=0.0012, Fig. 6B). The pyramidal cell layer was also significantly wider in bPAC-expressing CA1 compared to its contralateral counterpart (right CA1: 84.73 ± 2.23 µm, left CA1: 65.27 ± 0.94 µm, paired t-test: t=8.42, n=6, p=0.0004; Fig. 6C). Mice, which received the same viral vector, carrying just the reporter mCherry, had similar pyramidal cell layer widths in the left and right hippocampus (right CA1: 70.96 ± 3.53 µm, left CA1: 69.10 ± 3.17 µm, paired t-test: t=0.95, n=4, p=0.41; Fig. 6D). Cell dispersion, taken as the difference between right and left CA1 width, was the highest in bPAC-expressing CA1 (19.45 ± 2.31 µm), followed by PACK-expressing CA1 (10.98 ± 1.84 µm), and lacking in mCherry-expressing CA1 (1.86 ± 1.94 µm, one-way ANOVA: F(3,18)=12.51, p=0.0006, Tukey’s multiple comparisons test: PACK vs. bPAC p= 0.0302, PACK vs. mCherry p= 0.0397, bPAC vs. mCherry p= 0.0005; Fig. 6E). These findings led us to conclude that bPAC, and not the viral vector itself, is inducing the cell dispersion in the CA1 pyramidal cell layer.

**Figure 6.**
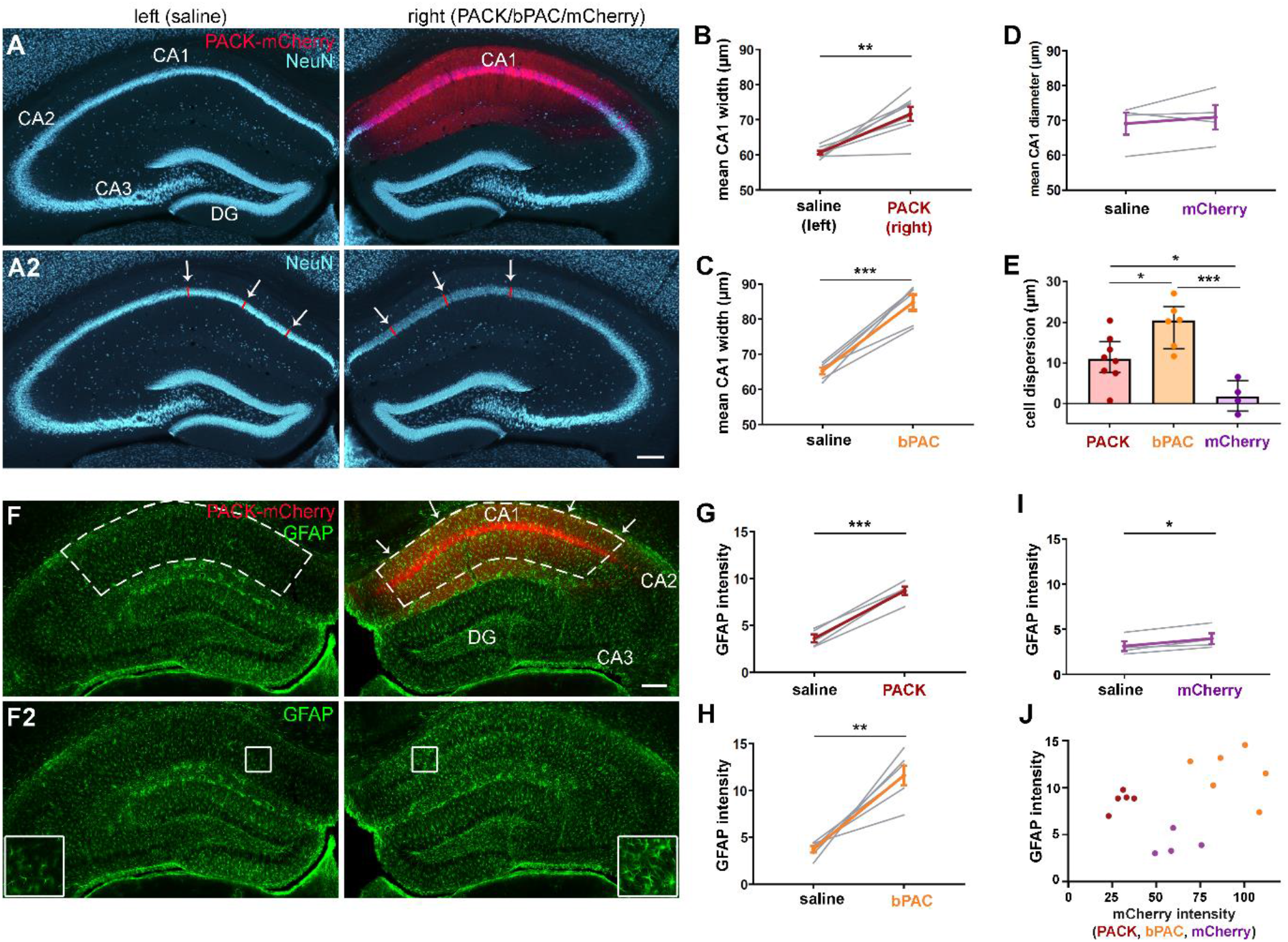
Cell dispersion and astrogliosis in PACK- and bPAC-expressing CA1. (A) Representative image of a NeuN-labeled hippocampal section with PACK-mCherry expression in the right CA1. The diameter of the CA1 pyramidal cell layer was measured at six positions (red lines and white arrows) in three hippocampal sections per animal. (B, C) The mean width of the CA1 pyramidal cell layer in (B) PACK-expressing (in red) and (C) in bPAC-expressing (in orange) hippocampus, was significantly increased compared to the contralateral saline-injected hippocampus (paired t-test). (D) In mCherry-expressing CA1, the pyramidal cell layer width was similar as in the contralateral CA1. (E) The cell dispersion (right-left CA1 width) was the strongest in bPAC mice (one-way ANOVA, Tukey’s post hoc).(F) In the representative image of a GFAP-stained section, GFAP labeling was visibly stronger in the PACK-expressing CA1. Mean grey value of GFAP labeling and mCherry expression was measured in the right dorsal CA1 including *str. oriens, str. pyramidale*, and *str. radiatum* (white dashed lines with arrows) in three sections per animal. GFAP intensity was measured at same positions in the left CA1 (white dashed lines). (G, H) GFAP labeling intensity was significantly higher in (G) PACK, (H) bPAC, and(I) mCherry-expressing CA1 compared to the left saline-injected side (paired t-test). (J) There was no clear correlation between GFAP labeling intensity and mCherry intensity, however PACK, bPAC, and mCherry mice formed three separate clusters with mCherry mice having the lowest GFAP labeling. *p<0.05, **p<0.01, ***p<0.001. Mean presented with SEM as error bars. Scale bars 200 µm.

GFAP labeling in hippocampal sections shows salient chronic astrogliosis in the PACK-expressing CA1 area (Fig. 6F, F2). The comparison of GFAP labeling in left and right hippocampi revealed strongly elevated GFAP intensity in the PACK-expressing CA1 (right CA1: 8.70 ± 0.46, left CA1: 3.62 ± 0.41, paired t-test: t=13.54, n=5, p= 0.0002; Fig. 6G). In bPAC mice, we also found notable astrogliosis in the right bPAC-expressing hippocampus (right CA1: 11.62 ± 1.03, left CA1: 3.73± 0.34, paired t-test: t=6.01, n=6, p=0.0018, Fig. 6H). In mCherry mice, the GFAP intensity was slightly but significantly elevated in the right hippocampus (right CA1: 3.98 ± 0.61, left CA1: 3.15± 0.53, paired t-test: t=4.12, n=4, p=0.026; Fig. 6I). Thus, it could be that either the viral vector or the presence of an electrode and an optic fiber contributed to the glial scarring in the right hippocampus. There was no significant correlation between the strength of mCherry-expression and GFAP intensity in PACK, bPAC, and mCherry mice (Pearson’s correlation: r= 0.35, p=0.2; Fig. 6J). However, the three groups formed separate clusters with: (1) mCherry mice having medium mCherry expression but the lowest GFAP intensity, (2) PACK mice having the lowest mCherry expression but medium GFAP intensity, and (3) bPAC mice having the strongest mCherry expression and the highest GFAP intensity. Taken together, it seems like bPAC expression is the main factor inducing chronic astrogliosis in PACK and bPAC mice, while the viral vector and hippocampal implantations might contribute additionally.

### PACK/bPAC expression in the contralateral hippocampus prevents seizure spread in chronically epileptic mice

To find out if the PACK silencer could be used to limit the spread of epileptiform activity, we targeted PACK to the CA1 principal cells, contralateral to ihpKA treatment. The ihpKA mouse model recapitulates the main pathological features of MTLE, i.e. spontaneous recurrent seizures associated with hippocampal sclerosis (Bouilleret et al., 1999). Most ihpKA mice have epileptiform activity that occurs in form of bursts originating in the seizure focus and propagating into the contralateral hippocampus (Meier et al., 2007; Janz et al., 2018; Paschen et al., 2020). Unexpectedly, all our PACK-injected ihpKA mice (n=5) were free of contralateral seizures already in baseline recordings before light-activation of PACK.

We detected the epileptiform bursts with high spike load using an automated algorithm (Heining et al., 2019) and quantified the burst ratio, the fraction of time spent in bursts, during the respective recording (Fig. 7). Although the epileptiform bursts occurred frequently in the ipsilateral hippocampus of PACK ihpKA mice (mean burst ratio in “pre”: 0.19 ± 0.02, “0.1 Hz”: 0.18 ± 0.02, “post”: 0.20 ± 0.02, n=5), the contralateral hippocampus was devoid of seizure activity (Fig. 7E-G). Similarly, in bPAC ihpKA mice, epileptiform bursts were detected in the ipsilateral (mean burst ratio in “pre”: 0.11 ± 0.03, “0.1 Hz”: 0.07 ± 0.03, “post”: 0.13 ± 0.05, n=3; Fig. 7I) but not the contralateral hippocampus (Fig. 7J). Neither the light-induced activation of bPAC nor the whole PACK construct had any effect on the burst ratios in the ipsilateral or contralateral hippocampus (Fig. 7E-J).

**Figure 7.**
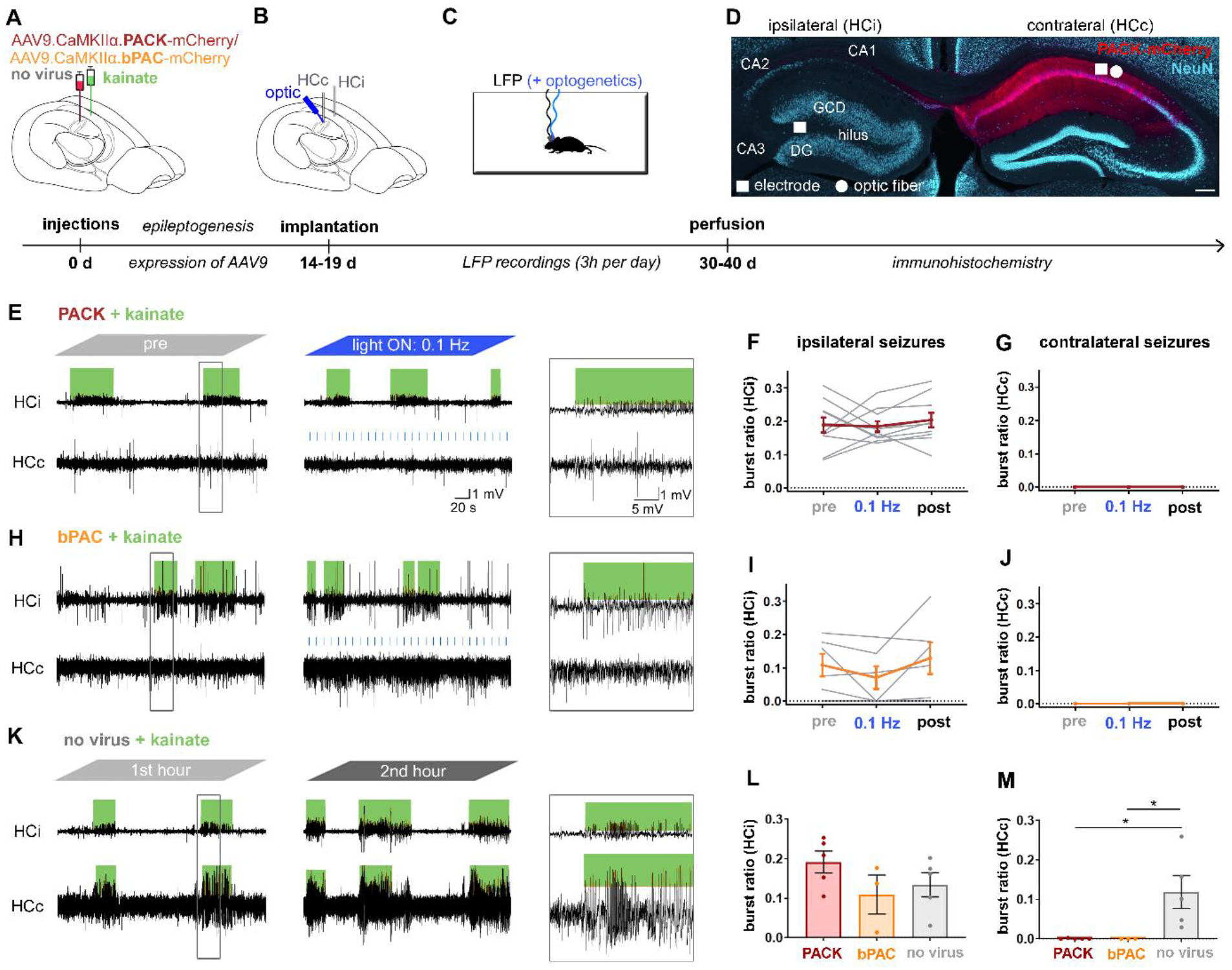
Chronically epileptic PACK and bPAC mice lack spontaneous seizures in the contralateral hippocampus. (A) Mice were injected with kainate in the left hippocampus and AAV9 carrying PACK, bPAC, or no virus in the contralateral hippocampus. (B) Two weeks later, wire electrodes were implanted into the kainate-injected ipsilateral hippocampus (HCi) and the virus-injected contralateral hippocampus (HCc). In PACK and bPAC mice, an optic fiber was implanted at a 30° angle adjacent to HCc. (C) Three-hour LFP recordings with and without optogenetic manipulations were performed as previously in healthy PACK/bPAC mice. ‘No virus’ mice were recorded only for three-hour reference recordings. (D) Representative section of an ihpKA PACK mouse showing hippocampal sclerosis in HCi with cell loss in CA1, CA3, and hilus regions as well as granule cell dispersion (GCD). PACK-mCherry was expressed in dorsal CA1 of HCc. Representative LFP snippets from (E) PACK-expressing (H) bPAC-expressing and (K) ‘no virus’ kainate-injected mice in the chronic phase of epilepsy. (E-M) Spontaneous epileptiform bursts (hypersynchronous spiking activity, marked in green) were detected by an automated algorithm (Heining et al., 2019), and quantified as burst ratio, a fraction of recording spent in bursts. (E-G) PACK and (H-J) bPAC mice had regularly occurring seizures in HCi but no propagation to HCc during pre-recordings as well as during and after illumination. (K) In contrast, mice without virus expression, frequently showed seizure propagation to HCc. (L) The burst ratio in HCi was similar in PACK, bPAC, and ‘no virus’ mice, (M) whereas in HCc, the burst ratio was significantly above zero in ‘no virus’ epileptic mice (33-36 days after ihpKA). One-way ANOVA with Dunnett’s multiple comparison test, *p<0.05. Mean presented with SEM as error bars.

We compared the burst ratios in PACK and bPAC mice to ‘no virus’ mice (n=4) in a three-hour recording 33-36 days after kainate to clarify whether the lack of contralateral seizures affects the seizure burden in the kainate-injected ipsilateral hippocampus (Fig. 7K-M). There was no significant difference in the ipsilateral burst ratios (PACK: 0.19 ± 0.03, bPAC: 0.11 ± 0.05, no virus: 0.13 ± 0.03, one-way ANOVA: F=1.53, n=3-5, p=0.26; Fig. 7L), whereas the mean contralateral burst ratio was evidently higher in ‘no virus’ mice than in PACK and bPAC mice (PACK: 0.00 ± 0.00, bPAC: 0.00 ± 0.00, no virus: 0.12 ± 0.04, one-way ANOVA: F=6.26, n=3-5, p= 0.017, Dunnett’s multiple comparisons test: PACK vs. no virus p=0.018, bPAC vs. no virus p=0.036; Fig. 7M). These results suggest that the dark activity of bPAC in CA1 pyramidal neurons prevents the spread of epileptiform activity to the bPAC-expressing areas. The activity of CA1 pyramidal cells is thus critical for seizure propagation into the contralateral hippocampus. Additionally, the absence of contralateral bursts does not affect the seizure burden in the sclerotic hippocampus.

## Discussion

The present study provides a detailed characterization of a novel two-component potassium channel-based silencer, PACK, and its long-term application in freely moving mice. We targeted the PACK construct to hippocampal CA1 with an AAV9.CaMKII vector, resulting in robust expression of PACK-mCherry in principal neurons of this area. We report that PACK is a suitable silencer to reduce the activity of hippocampal neurons *in vivo* by applying short blue light pulses via an implanted optic fiber. Previously, Bernal Sierra et al. (2018) demonstrated in acute hippocampal slices that shining 5 ms blue light pulses at 0.05 Hz onto PACK-expressing CA1 cells was sufficient to abolish spiking elicited by current injections. In our freely moving mice, applying 10 ms light pulses at 0.05 Hz provided an unstable lowering of the LFP magnitude with periods of recovery. However, with an increased illumination frequency of 0.1 Hz, neuronal activity was persistently reduced throughout the 50-minute light ON phase. This is a significant improvement compared to microbial chloride and proton pumps, which require continuous illumination and already exhibit declining photocurrent amplitudes within one minute (Mattis et al., 2012; Wiegert et al., 2017).

Regarding oscillatory activity in CA1, light-induced PACK activation in CA1 pyramidal cells mainly reduced the power of gamma and beta oscillations. Hippocampal gamma rhythm depends on the firing of pyramidal cells and their synchronized dendritic and perisomatic inhibitory input, which originate from local somatostatin- and parvalbumin-positive interneurons, respectively (Tukker et al., 2007; Antonoudiou et al., 2020). Constant reduction of pyramidal cell activity consequently decreases local interneuron activity (Gridchyn et al., 2020), which would explain the lower power of gamma oscillations during PACK-mediated inhibition. The beta power was probably reduced for the same reason, as the activity of local CA1 interneurons is also crucial for beta oscillations (Rangel et al., 2016). PACK-mediated inhibition affected theta oscillations to a much lesser extent, probably while theta rhythm is predominantly driven by GABAergic and cholinergic inputs from the medial septum while depending less on local excitation and inhibition (Hangya et al., 2009; Fuhrmann et al., 2015; Müller and Remy, 2018).

Using soluble bPAC as the light-sensitive domain in the PACK tool is favorable to ensure high light sensitivity (Beck et al., 2018; Bernal Sierra et al., 2018). However, this advantage seems to have come with dark activity, which is probably the reason for the side effects we observed in PACK- and bPAC-expressing hippocampi *in vivo*. The harmful effects of chronic bPAC-expression in the hippocampal CA1 area included pyramidal cell layer widening, chronic astrogliosis, and spontaneous generalized seizures. Mice that received the AAV9 vector carrying only mCherry under the CaMKII promoter did not show these side effects, except for astrogliosis, which could be due to implantation-related scarring. Based on these findings, we conclude that the viral vector and the fluorescent marker alone were not detrimental. As bPAC mice tended to have higher expression of bPAC-mCherry, more prominent CA1 dispersion, stronger astrogliosis, and higher occurrence of generalized seizures than PACK mice, chronic cAMP elevation may be the underlying cause of these adverse effects.

The second messenger molecule cAMP has several molecular targets in neurons, including protein kinase A (PKA), exchange protein activated by cAMP (Epac), and cAMP-gated ion channels. Through these pathways, cAMP regulates fundamental physiological processes such as growth, metabolism, migration, apoptosis, gene transcription, neurotransmission, and plasticity (for review, see Nguyen & Woo, 2003; Cheng et al., 2008; Antoni, 2012). Altered gene expression by activation of cAMP-responsive element (CRE) could explain histological changes and seizure activity in bPAC mice as the CRE-transcriptional pathway is involved in acute and chronic phases of epilepsy (Huang et al., 1994; Hansen et al., 2014; Choi et al., 2016; Conte et al., 2020). The nuclear distribution element-like 1 (NDEL1) protein, transcribed in an activity-dependent manner by the cAMP response element-binding (CREB) protein, has been hypothesized to contribute to pathophysiological alterations of MTLE. NDEL1 is robustly and persistently elevated in the mouse hippocampus, including CA1 neurons, after pilocarpine-induced *status epilepticus* (Choi et al., 2016; Zhu et al., 2020). A conditional knockout of the NDEL1 gene in CA1 of adolescent mice leads to dispersion and increased intrinsic excitability of CA1 pyramidal cells (Jiang et al., 2016), resembling what we observed in our bPAC mice.

Light-activation of bPAC *in vivo* strongly increased LFP magnitude and the oscillatory power of beta and gamma in the CA1 region. The neuronal activity dropped immediately after the illumination phase, suggesting a temporary depolarizing effect via a cyclic nucleotide-gated channel or induction of short-term potentiation. Local cAMP elevation induces presynaptic potentiation by promoting the accumulation of calcium channels close to release sites, thus increasing the release probability (Midorikawa and Sakaba, 2017; Vaden et al., 2019; Fukaya et al., 2021). Presynaptic elevation of cAMP via light-activation of synapse-targeted bPAC (synaptoPAC) was sufficient to trigger potentiation at the mossy fiber-CA3 synapse but not in the CA3-CA1 synapse (Oldani et al., 2021). Although cAMP-mediated presynaptic potentiation is not exhibited by all hippocampal synapses, it is present in the CA1-subicular bursting neuron synapse (Wozny et al., 2008). Therefore, temporarily increased transmission at the CA1-subiculum synapses might have contributed to light-induced hyperactivity in bPAC-expressing CA1 *in vivo*. Alternatively, cAMP binding on hyperpolarization-activated cyclic nucleotide-gated (HCN) cation channels might have increased pyramidal cell excitability because (1) HCN channels are abundantly expressed in hippocampal neurons, and (2) the inward currents through HCN channels depolarize the membrane (Santoro et al., 2000; Lörincz et al., 2002; He et al., 2014). Furthermore, HCN channels are thought to initiate rhythmic firing (Nolan et al., 2004; He et al., 2014), which could theoretically explain elevated beta and gamma oscillations during light-induced bPAC activation *in vivo*. Future experiments with probes or tetrodes could potentially clarify the mechanism of bPAC-mediated excitation and PACK-mediated inhibition at a single-unit level.

In the last part of our study, we applied the PACK silencer in chronically epileptic ihpKA mice, which usually exhibit spontaneous seizures in both hippocampi (Meier et al., 2007; Janz et al., 2018; Paschen et al., 2020). We targeted PACK to the contralateral hippocampus (opposite to kainate injection) to determine if we can interfere with seizure spread between the two hippocampi. To our surprise, there was no propagation of seizure activity from the kainate-injected hippocampus to the PACK-expressing contralateral side, even before light-activation of PACK. We saw the same in bPAC mice, which were also lacking contralateral seizures already in the baseline recordings. Surprisingly, prolonged bPAC expression in saline-injected mice induced generalized seizures, whereas it prevented seizure spread in kainate-injected mice. Hypothetically, bPAC-dependent cAMP elevation and subsequent HCN channel activation could explain contradictory findings regarding epileptogenicity, since HCN channels have also been found pro- and anti-epileptic (Bender et al., 2003; Lewis and Chetkovich, 2011; Noam et al., 2011; Stegen et al., 2012). Future work comparing gene expression and hippocampal slice electrophysiology in saline- and kainate-treated mice with and without bPAC-expression would be needed to address the mechanism of cAMP-associated seizure induction and prevention.

Our results suggest that the expression of bPAC affects the physiology of hippocampal principal cells in the absence of blue light. Functional dark activity of soluble bPAC has also been reported by others, who utilized a fluorescent PKA-sensor to detect intracellular cAMP levels in bPAC-expressing CA1 cells in organotypic hippocampal cultures (personal communication with O. Constantin, group of Prof. Oertner). For reduced dark activity, the soluble bPAC from the *Beggiatoa* bacterium could be replaced by another photoactivated adenylyl cyclase (PAC). There are several PACs available from other microorganisms, which have lower light sensitivity but still provide sufficient potassium current when coupled to SthK (Bernal Sierra et al., 2018). Alternatively, a red-shifted bPAC with reduced dark activity could be used (Stierl et al., 2014). Another approach would be targeting a PAC to the cell membrane. For example, membrane-bound guanylyl cyclase rhodopsins, which have mutated to be adenylyl cyclases, virtually lack dark activity (Scheib et al., 2018; Henß et al., 2021). In these approaches, the possibility of SthK channel activation by intrinsic cAMP remains. Accordingly, expression of SthK without bPAC in body wall muscle cells of *C*.*elegans* was already sufficient to see a behavioral change resulting from muscle hyperpolarization (Henß et al., 2021). This problem could be overcome by engineering SthK variants with mutations in the cAMP binding site, resulting in a channel with reduced cAMP affinity.

In conclusion, we showed in awake mice that the potassium channel-based optogenetic silencer PACK reliably reduces hippocampal neuronal activity in a light-dependent manner. In contrast to other optogenetic inhibitors, PACK requires only short light pulses at a low frequency to achieve prolonged reduction of neuronal activity. A disadvantage of PACK is its light-active component, bPAC, since it elicits side effects *in vivo*, which are presumably related to its dark activity. In the mouse model of MTLE, the light-independent effects of bPAC prevented the spread of spontaneous epileptiform activity from the seizure focus to the contralateral bPAC-expressing CA1 region. Our study underlines that the PACK tool is a potent optogenetic inhibitor *in vivo* but refinement of its light-sensitive domain is required to avoid dark activity and related side effects.

## Acknowledgements

We are grateful to the Viral Core Facility, Charité – Universitätsmedizin Berlin, Germany for providing the viral constructs. We thank Prof. Peter Hegemann and Philipp Janz for intellectual input, Andrea Djie-Maletz for excellent technical assistance, Jessica Link for proofreading, and Oana Constantin for unpublished information about the photoactivated adenylyl cyclase, bPAC.

This study was supported by the German Research Foundation (HA 1443/11-1 to CAH, SPP1926 to YABS) and by the BrainLinks-BrainTools Center, which is funded by the Federal Ministry of Economics, Science and Arts of Baden-Württemberg within the sustainability program for projects of the Excellence Initiative II.

## Supplementary data

**Supplementary figure 1.**
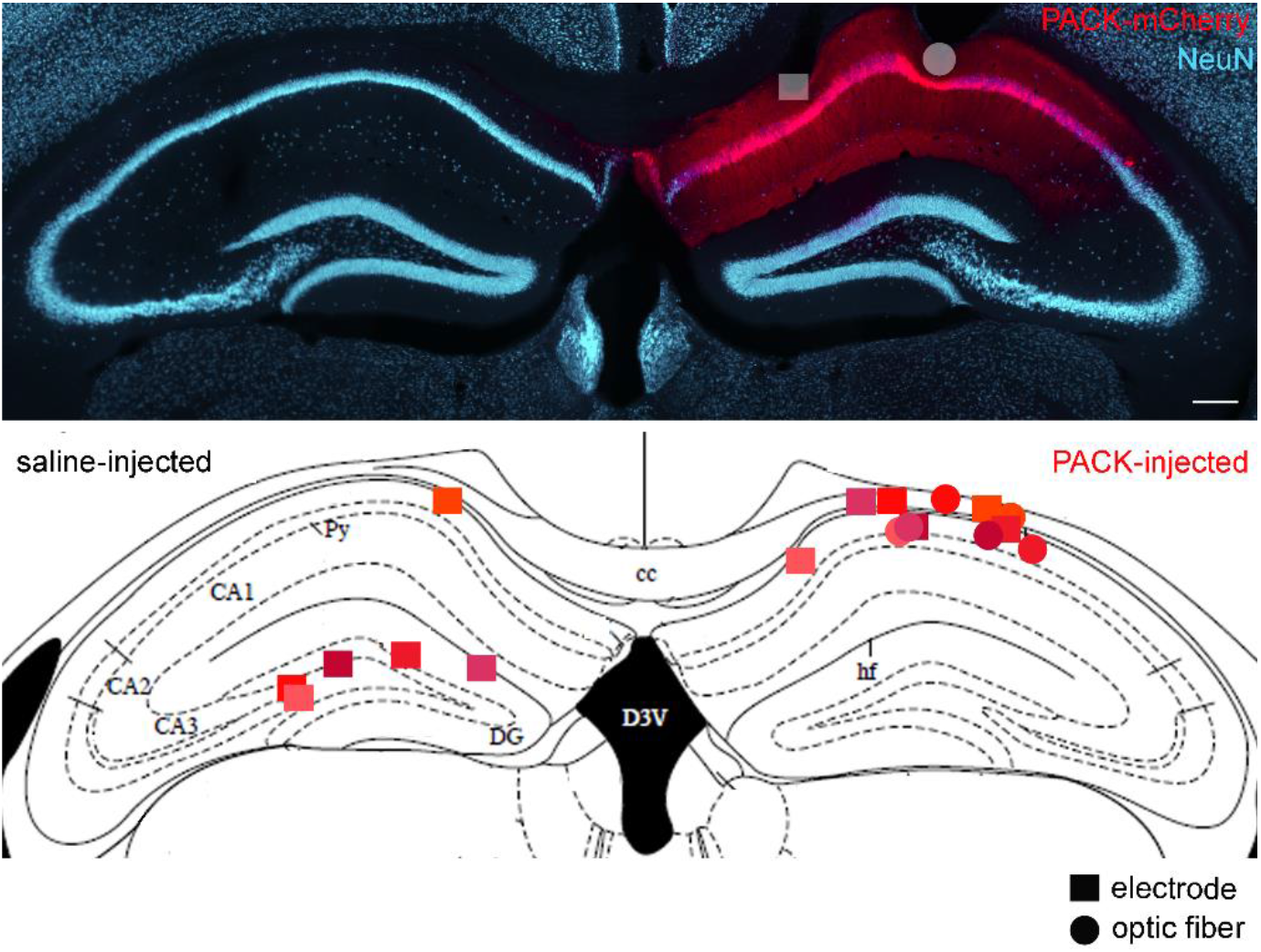
Electrode and optic fiber positions in dorsal hippocampi of PACK mice. (A) A representative NeuN-labeled hippocampal section, where the electrode (square) and optic fiber (circle) positions are visible in PACK-expressing right hippocampus. (B) Electrode and optic fiber positions in saline-injected and PACK-injected hippocampi of mice that were included in the LFP analysis (n=6, in different shades of red).

**Supplementary figure 2.**
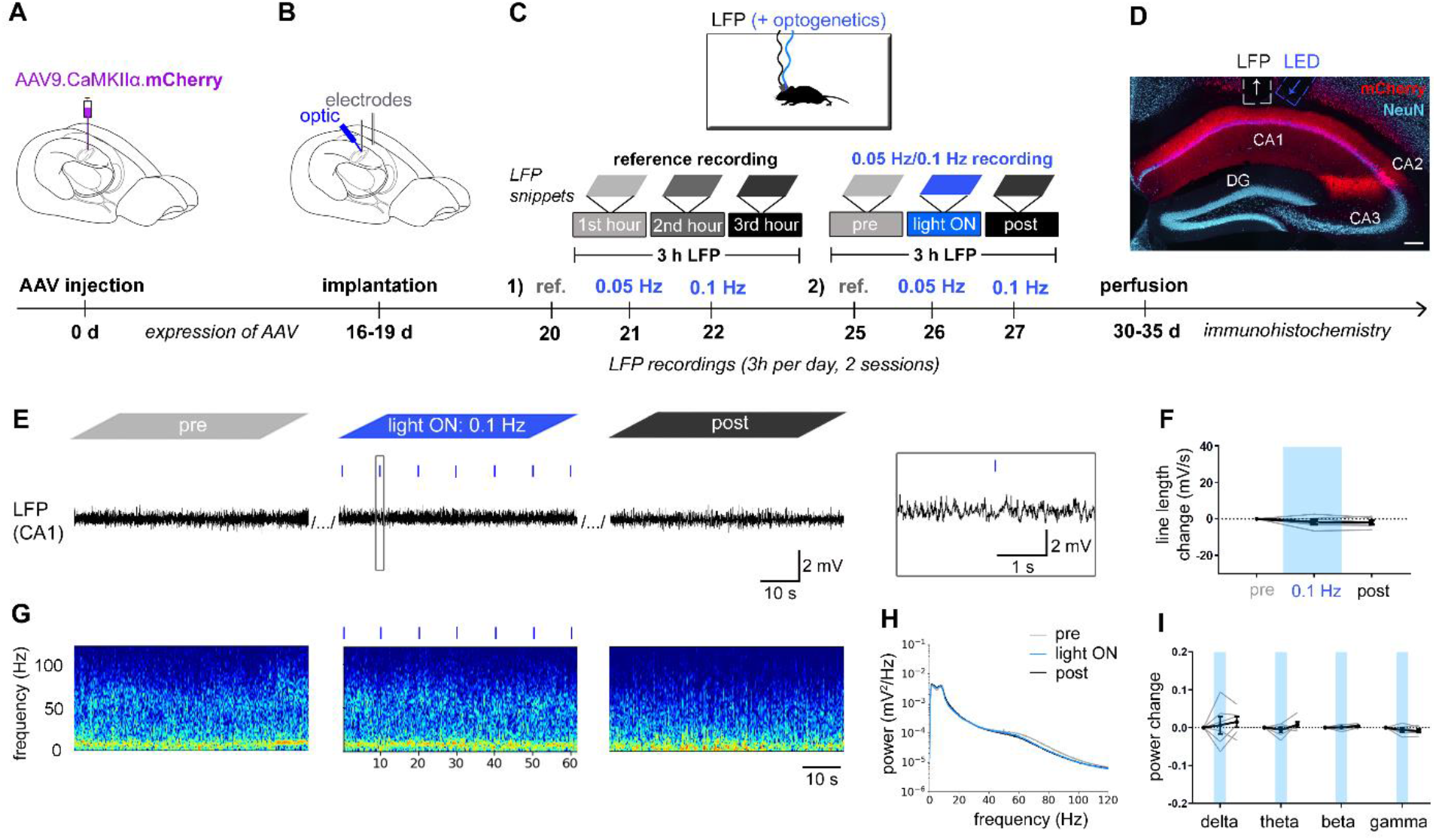
Light application in mCherry mice does not alter neuronal activity in CA1. Experimental design. We targeted mCherry to excitatory neurons in the CA1 area using the AAV9 vector. (B) Implantation, (C) recordings and (D) histological analysis were performed like in PACK mice. Scale bar 200 µm. (E) Representative LFP snippets of a 0.1 Hz recording show no response to the light application. (F) Line length was not changed by 0.1 Hz illumination. (G) The spectrograms, taken from the same time windows as LFP snippets, were unaltered during 0.1 Hz illumination. (H) The average PSD is nearly overlapping during pre-, light ON, and post-recordings, suggesting the light application did not affect the oscillations in the CA1 region. (I) The power of delta, theta, beta, and gamma oscillations remained the same across the three recording hours, not being altered by the illumination (shown in blue).

## References

Alfonsa H, Merricks EM, Codadu NK, Cunningham MO, Deisseroth K, Racca C, Trevelyan AJ (2015) The contribution of raised intraneuronal chloride to epileptic network activity. J Neurosci 35:7715–7726.

Antoni FA (2012) New paradigms in cAMP signalling. Mol Cell Endocrinol 353:3–9.

Antonoudiou P, Tan YL, Kontou G, Louise Upton A, Mann EO (2020) Parvalbumin and somatostatin interneurons contribute to the generation of hippocampal gamma oscillations. J Neurosci 40:7668–7687.

Banghart M, Borges K, Isacoff E, Trauner D, Kramer RH (2004) Light-activated ion channels for remote control of neuronal firing. Nat Neurosci 7:1381–1386.

Beck S, Yu-Strzelczyk J, Pauls D, Constantin OM, Gee CE, Ehmann N, Kittel RJ, Nagel G, Gao S (2018) Synthetic light-activated ion channels for optogenetic activation and inhibition. Front Neurosci 12:643.

Bender RA, Soleymani S V., Brewster AL, Nguyen ST, Beck H, Mathern GW, Baram TZ (2003) Enhanced expression of a specific hyperpolarization-activated cyclic nucleotide-gated cation channel (HCN) in surviving dentate gyrus granule cells of human and experimental epileptic hippocampus. J Neurosci 23:6826–6836.

Bernal Sierra YA, Rost BR, Pofahl M, Fernandes AM, Kopton RA, Moser S, Holtkamp D, Masala N, Beed P, Tukker JJ, Oldani S, Bönigk W, Kohl P, Baier H, Schneider-Warme F, Hegemann P, Beck H, Seifert R, Schmitz D (2018) Potassium channel-based optogenetic silencing. Nat Commun 9:4611.

Bouilleret V, Ridoux V, Depaulis A, Marescaux C, Nehlig A, Le Gal La Salle G (1999) Recurrent seizures and hippocampal sclerosis following intrahippocampal kainate injection in adult mice: Electroencephalography, histopathology and synaptic reorganization similar to mesial temporal lobe epilepsy. Neuroscience 89:717–729.

Bragin A, Jando G, Nadasdy Z, Hetke J, Wise K, Buzsaki G (1995) Gamma (40-100 Hz) oscillation in the hippocampus of the behaving rat. J Neurosci 15:47–60.

Brams M, Kusch J, Spurny R, Benndorf K, Ulens C (2014) Family of prokaryote cyclic nucleotide-modulated ion channels. Proc Natl Acad Sci USA 111:7855–7860.

Buzsáki G, Buhl DL, Harris KD, Csicsvari J, Czéh B, Morozov A (2003) Hippocampal network patterns of activity in the mouse. Neuroscience 116:201–211.

Cabib S, Algeri S, Perego C, Puglisi-Allegra S (1990) Behavioral and biochemical changes monitored in two inbred strains of mice during exploration of an unfamiliar environment. Physiol Behav 47:749–753.

Cheng X, Ji Z, Tsalkova T, Mei F (2008) Epac and PKA: A tale of two intracellular cAMP receptors. Acta Biochim Biophys Sin (Shanghai) 40:651–662.

Choi YS, Lee B, Hansen KF, Aten S, Horning P, Wheaton KL, Impey S, Hoyt KR, Obrietan K (2016) Status epilepticus stimulates NDEL1 expression via the CREB/CRE pathway in the adult mouse brain. Neuroscience 331:1–12.

Colgin LL (2016) Rhythms of the hippocampal network. Nat Rev Neurosci 17:239– 249.

Conte G, Parras A, Alves M, Ollà I, De Diego-Garcia L, Beamer E, Alalqam R, Ocampo A, Mendez R, Henshall DC, Lucas JJ, Engel T (2020) High concordance between hippocampal transcriptome of the mouse intra-amygdala kainic acid model and human temporal lobe epilepsy. Epilepsia 61:2795–2810.

Cosentino C, Alberio L, Gazzarrini S, Aquila M, Romano E, Cermenati S, Zuccolini P, Petersen J, Beltrame M, Etten JL Van, Christie JM, Thiel G, Moroni A (2015) Optogenetics. Engineering of a light-gated potassium channel. Science 348:707–710.

Engel J.J (2001) Mesial temporal lobe epilepsy: What have we learned? Neuroscientist 7:340–352.

Esteller R, Echauz J, Tcheng T (2004) Comparison of line length feature before and after brain electrical stimulation in epileptic patients. Conf Proc IEEE Eng Med: 707–710.

Feng G, Mellor RH, Bernstein M, Keller-Peck C, Nguyen QT, Wallace M, Nerbonne JM, Lichtman JW, Sanes JR (2000) Imaging neuronal subsets in transgenic mice expressing multiple spectral variants of GFP. Neuron 28:41–51.

Fuhrmann F, Justus D, Sosulina L, Kaneko H, Beutel T, Friedrichs D, Schoch S, Schwarz MK, Fuhrmann M, Remy S (2015) Locomotion, theta oscillations, and the speed-correlated firing of hippocampal neurons are controlled by a medial septal glutamatergic circuit. Neuron 86:1253–1264.

Fukaya R, Maglione M, Sigrist SJ, Sakaba T (2021) Rapid Ca2+ channel accumulation contributes to cAMP-mediated increase in transmission at hippocampal mossy fiber synapses. Proc Natl Acad Sci USA 118: e2016754118.

Gloor P, Salanova V, Olivier A, Quesney LF (1993) The human dorsal hippocampal commissure: An anatomically identifiable and functional pathway. Brain 116:1249–1273.

Gridchyn I, Schoenenberger P, O’neill J, Csicsvari J (2020) Optogenetic inhibition-mediated activity-dependent modification of CA1 pyramidal-interneuron connections during behavior. Elife 9:e61106.

Gruart A, Benito E, Delgado-García JM, Barco A (2012) Enhanced cAMP response element-binding protein activity increases neuronal excitability, hippocampal long-term potentiation, and classical eyeblink conditioning in alert behaving mice. J Neurosci 32:17431–17441.

Hangya B, Borhegyi Z, Szilágyi N, Freund TF, Varga V (2009) GABAergic neurons of the medial septum lead the hippocampal network during theta activity. J Neurosci 29:8094–8102.

Hansen KF, Sakamoto K, Pelz C, Impey S, Obrietan K (2014) Profiling status epilepticus-induced changes in hippocampal RNA expression using high-throughput RNA sequencing. Sci Rep 4:6930.

Häussler U, Bielefeld L, Froriep UP, Wolfart J, Haas CA (2012) Septotemporal position in the hippocampal formation determines epileptic and neurogenic activity in temporal lobe epilepsy. Cereb Cortex 22:26–36.

He C, Chen F, Li B, Hu Z (2014) Neurophysiology of HCN channels: From cellular functions to multiple regulations. Prog Neurobiol 112:1–23.

Heining K, Kilias A, Janz P, Häussler U, Kumar A, Haas CA, Egert U (2019) Bursts with high and low load of epileptiform spikes show context-dependent correlations in epileptic mice. eNeuro 6:ENEURO.0299-18.2019.

Heinrich C, Nitta N, Flubacher A, Müller M, Fahrner A, Kirsch M, Freiman T, Suzuki F, Depaulis A, Frotscher M, Haas CA (2006) Reelin deficiency and displacement of mature neurons, but not neurogenesis, underlie the formation of granule cell dispersion in the epileptic hippocampus. J Neurosci 26:4701–4713.

Henß T, Nagpal J, Gao S, Scheib U, Pieragnolo A, Hirschhäuser A, Schneider-Warme F, Hegemann P, Nagel G, Gottschalk A (2021) Optogenetic tools for manipulation of cyclic nucleotides functionally coupled to cyclic nucleotide-gated channels. Br J Pharmacol: 10.1111/bph.15445.

Hinman JR, Penley SC, Long LL, MA Escabí, Chrobak JJ (2011) Septotemporal variation in dynamics of theta: Speed and habituation. J Neurophysiol 105:2675– 2686.

Huang YY, Li XC, Kandel ER (1994) cAMP contributes to mossy fiber LTP by initiating both a covalently mediated early phase and macromolecular synthesis-dependent late phase. Cell 79:69–79.

Janovjak H, Szobota S, Wyart C, Trauner D, Isacoff EY (2010) A light-gated, potassium-selective glutamate receptor for the optical inhibition of neuronal firing. Nat Neurosci 13:1027–1032.

Janz P, Hauser P, Heining K, Nestel S, Kirsch M, Egert U, Haas CA (2018) Position- and time-dependent arc expression links neuronal activity to synaptic plasticity during epileptogenesis. Front Cell Neurosci 12:244.

Janz P, Savanthrapadian S, Häussler U, Kilias A, Nestel S, Kretz O, Kirsch M, Bartos M, Egert U, Haas CA (2017) Synaptic Remodeling of Entorhinal Input Contributes to an Aberrant Hippocampal Network in Temporal Lobe Epilepsy. Cereb Cortex 27:2348–2364.

Jiang Y, Gavrilovici C, Chansard M, Liu RH, Kiroski I, Parsons K, Park SK, Teskey GC, Rho JM, Nguyen MD (2016) Ndel1 and reelin maintain postnatal CA1 hippocampus integrity. J Neurosci 36:6538–6552.

Kang JY, Kawaguchi D, Coin I, Xiang Z, DDM O’Leary, Slesinger PA, Wang L (2013) In vivo expression of a light-activatable potassium channel using unnatural amino acids. Neuron 80:358–370.

Kwan P, Sander JW (2004) The natural history of epilepsy: An epidemioloqical view. J Neurol Neurosurg Psychiatry 75:1376–1381.

Lewis AS, Chetkovich DM (2011) HCN channels in behavior and neurological disease: Too hyper or not active enough? Mol Cell Neurosci 46:357–367.

Lörincz A, Notomi T, Tamás G, Shigemoto R, Nusser Z (2002) Polarized and compartment-dependent distribution of HCN1 in pyramidal cell dendrites. Nat Neurosci 5:1185–1193.

Mahn M, Prigge M, Ron S, Levy R, Yizhar O (2016) Biophysical constraints of optogenetic inhibition at presynaptic terminals. Nat Neurosci 19:554–556.

Mattis J, Tye KM, Ferenczi EA, Ramakrishnan C, O’Shea DJ, Prakash R, Gunaydin LA, Hyun M, Fenno LE, Gradinaru V, Yizhar O, Deisseroth K (2012) Principles for applying optogenetic tools derived from direct comparative analysis of microbial opsins. Nat Methods 9:159–172.

Meier R, Häussler U, Aertsen A, Deransart C, Depaulis A, Egert U (2007) Short-term changes in bilateral hippocampal coherence precede epileptiform events. Neuroimage 38:138–149.

Midorikawa M, Sakaba T (2017) Kinetics of Releasable Synaptic Vesicles and Their Plastic Changes at Hippocampal Mossy Fiber Synapses. Neuron 96(5):1033-1040.e3.

Mintzer S, Cendes F, Soss J, Andermann F, Engel J, Dubeau F, Olivier A, Fried I (2004) Unilateral hippocampal sclerosis with contralateral temporal scalp ictal onset. Epilepsia 45:792–802.

Müller C, Remy S (2018) Septo – hippocampal interaction. Cell Tissue Res 373:565– 575.

Nguyen P V., Woo NH (2003) Regulation of hippocampal synaptic plasticity by cyclic AMP-dependent protein kinases. Prog Neurobiol 71:401–437.

Noam Y, Bernard C, Baram TZ (2011) Towards an integrated view of HCN channel role in epilepsy. Curr Opin Neurobiol 21:873–879.

Nolan M, Malleret G, Dudman J, Buhl D, Santoro B, Gibbs E, Vronskaya S, Buzsaki G, Siegelbaum S, Kandel E (2004) A behavioral role for dendritic integration: HCN1 channels constrain spatial memory and plasticity at inputs to distal dendrites of CA1 pyramidal neurons. Cell 119:719–732.

Oldani S, Moreno-Velasquez L, Faiss L, Stumpf A, Rosenmund C, Schmitz D, Rost BR (2021) SynaptoPAC, an optogenetic tool for induction of presynaptic plasticity. J Neurochem 156:324–336.

Owen SF, Liu MH, Kreitzer AC (2019) Thermal constraints on in vivo optogenetic manipulations. Nat Neurosci 22:1061–1065.

Paschen E, Elgueta C, Heining K, Vieira DM, Kleis P, Orcinha C, Häussler U, Bartos M, Egert U, Janz P, Haas CA (2020) Hippocampal low-frequency stimulation prevents seizure generation in a mouse model of mesial temporal lobe epilepsy. Elife 9:e54518.

Popovic L, Vojvodic N, Ristic AJ, Bascarevic V, Sokic D, Kostic VS (2012) Ictal dystonia and secondary generalization in temporal lobe seizures: A video-EEG study. Epilepsy Behav 25:501–504.

Racine RJ (1972) Modification of seizure activity by electrical stimulation. II. Motor seizure. Electroencephalogr Clin Neurophysiol 32:281–294.

Raimondo J V., Kay L, Ellender TJ, Akerman CJ (2012) Optogenetic silencing strategies differ in their effects on inhibitory synaptic transmission. Nat Neurosci 15:1102–1104.

Rangel LM, Rueckemann JW, Riviere PD, Keefe KR, Porter BS, Heimbuch IS, Budlong CH, Eichenbaum H (2016) Rhythmic coordination of hippocampal neurons during associative memory processing. Elife 5:1–24.

Riban V, Bouilleret V, Pham-Lê BT, Fritschy JM, Marescaux C, Depaulis A (2002) Evolution of hippocampal epileptic activity during the development of hippocampal sclerosis in a mouse model of temporal lobe epilepsy. Neuroscience 112:101–111.

Santoro B, Chen S, Lüthi A, Pavlidis P, Shumyatsky GP, Tibbs GR, Siegelbaum SA (2000) Molecular and functional heterogeneity of hyperpolarization-activated pacemaker channels in the mouse CNS. J Neurosci 20:5264–5275.

Scheib U, Broser M, Constantin OM, Yang S, Gao S, Mukherjee S, Stehfest K, Nagel G, Gee CE, Hegemann P (2018) Rhodopsin-cyclases for photocontrol of cGMP/cAMP and 2.3 Å structure of the adenylyl cyclase domain. Nat Commun 9:707–710.

Sørensen AT, Ledri M, Melis M, Ledri LN, Andersson M, Kokaia M (2017) Altered chloride homeostasis decreases the action potential threshold and increases hyperexcitability in hippocampal neurons. eNeuro 4:ENEURO.0172-17.2017.

Stegen M, Kirchheim F, Hanuschkin A, Staszewski O, Veh RW, Wolfart J (2012) Adaptive intrinsic plasticity in human dentate gyrus granule cells during temporal lobe epilepsy. Cereb Cortex 22:2087–2101.

Stierl M, Stumpf P, Udwari D, Gueta R, Hagedorn R, Losi A, Gärtner W, Petereit L, Efetova M, Schwarzel M, Oertner TG, Nagel G, Hegemann P (2011) Light modulation of cellular cAMP by a small bacterial photoactivated adenylyl cyclase, bPAC, of the soil bacterium Beggiatoa. J Biol Chem 286:1181–1188.

Tukker JJ, Fuentealba P, Hartwich K, Somogyi P, Klausberger T (2007) Cell type-specific tuning of hippocampal interneuron firing during gamma oscillations in vivo. J Neurosci 27:8184–8189.

Tulke S, Haas CA, Häussler U (2019) Expression of brain-derived neurotrophic factor and structural plasticity in the dentate gyrus and CA2 region correlate with epileptiform activity. Epilepsia 60:1234–1247.

Twele F, Schidlitzki A, Töllner K, Löscher W (2017) The intrahippocampal kainate mouse model of mesial temporal lobe epilepsy: Lack of electrographic seizure-like events in sham controls. Epilepsia Open 2:180–187.

Vaden JH, Banumurthy G, Gusarevich ES, Overstreet-Wadiche L, Wadiche JI (2019) The readily-releasable pool dynamically regulates multivesicular release. Elife 8:8:e47434.

Weisskopf MG, Castillo PE, Zalutsky RA, Nicoll RA (1994) Mediation of Hippocampal Mossy Fiber Long-Term Potentiation by Cyclic AMP. Science 265:1878–1882.

Wiegert JS, Mahn M, Prigge M, Printz Y, Yizhar O (2017) Silencing Neurons: Tools, Applications, and Experimental Constraints. Neuron 95:504–529.

Wozny C, Maier N, Fidzinski P, Breustedt J, Behr J, Schmitz D (2008) Differential cAMP signaling at hippocampal output synapses. J Neurosci 28:14358–14362.

Zhu L, Dai S, Lu D, Xu P, Chen L, Han Y, Zhong L, Chang L, Wu Q (2020) Role of NDEL1 and VEGF/VEGFR-2 in mouse hippocampus after status epilepticus. ASN Neuro 12:1759091420926836.

